# Tongue immune compartment analysis reveals spatial macrophage heterogeneity

**DOI:** 10.1101/2022.02.09.479699

**Authors:** Ekaterini Maria Lyras, Karin Zimmermann, Lisa Katharina Wagner, Dorothea Dörr, Cornelius Fischer, Steffen Jung, Simon Yona, Avi-Hai Hovav, Werner Stenzel, Steffen Dommerich, Thomas Conrad, Achim Leutz, Alexander Mildner

## Abstract

The tongue is a unique muscular organ situated in the oral cavity where it is involved in taste sensation, mastication and articulation. As a barrier organ, which is constantly exposed to environmental pathogens, the tongue is expected to host an immune cell network ensuring local immune defence. However, the composition and the transcriptional landscape of the tongue immune system are currently not completely defined. Here we characterised the tissue-resident immune compartment of the murine tongue during development, health and disease, combining single cell RNA-sequencing with *in situ* immunophenotyping. We identified distinct local immune cell populations and described two specific subsets of tongue-resident macrophages occupying discrete anatomical niches. *Cx3cr1*^+^ macrophages were located specifically in the highly innervated lamina propria beneath the tongue epidermis and at times in close proximity to fungiform papillae. *Folr2*^+^ macrophages were detected in deeper muscular tissue. The two macrophage subsets originate from a common proliferative precursor during early postnatal development and responded differently to systemic LPS *in vivo*. Our description of the under-investigated tongue immune system sets a starting point to facilitate research on tongue immune-physiology and pathology including cancer and taste disorders.

## Introduction

The tongue is a highly innervated muscular organ with functions in articulation, mastication and taste perception. Located at the entrance of the gastrointestinal tract, the tongue is constantly exposed to dietary and airborne antigens and therefore acts as a first-line immune organ ^1^. Moreover, taste sensation plays a critical role in avoidance of spoiled food and beverages. Accordingly, a tongue-resident immune network would be expected with roles in immune defense, tissue remodeling and tongue homeostasis. However, in the immunological context, the tongue is an understudied organ and the composition of tongue immune cells and their transcriptional status is largely unknown. Here, we define the immune cell landscape of the tongue with a specific focus on mononuclear phagocytes, e.g. tissue-resident macrophages (TRM).

Until now, the characterization of the mononuclear phagocyte compartment of the tongue mainly focused on Langerhans cells that were first described in the mouse epithelium forty years ago ^2^ and are identified in humans by their exclusive CD1a immunoreactivity ^3,4^. The characterization of sub-epithelial macrophage subsets is much more enigmatic and so far depended on the histological examination of a few membrane markers ^5,6^. For example, human CD163^+^ macrophages can be found in subepithelial areas of the tongue ^6^, a localization that they share with CD11c^+^ “dendritic” cells ^7^. Relying on single markers such as CD11c to identify cells of the “dendritic” cell lineage is problematic since various additional cell types, including monocytes, macrophages and lymphocytes, can express CD11c ^8^. Besides histological examinations of mononuclear phagocytes in the healthy tongue, the macrophage involvement in various pathological settings has also been studied. Tongue Langerhans cells were for instance shown to be critically involved in IL17-dependent antifungal immunity in the oral mucosa ^9^, play an important role in T cell priming during squamous cell carcinoma development ^10^ and are depleted in patients with advanced-stage acquired immune deficiency syndrome ^11^. Furthermore, an increase of activated ED1^+^ tongue macrophages was observed in systemic inflammation in rats ^12^. The recent COVID-19 pandemic has also suggested a potential link of viral infections with tongue immunity, with loss of taste being one of the hallmark symptoms ^13^. However, the lack of knowledge of the tongue immune cell compartment in physiology, hampers our understanding of tongue immune responses following pathogen challenge. Therefore, an unbiased characterization of the tongue immune cells is critical to classify and evaluate tissue-resident cell subsets, e.g. macrophage dynamics during tongue development and pathologies.

To this end, we profiled the tongue-resident CD45^+^ hematopoietic cell compartment by single cell RNA-sequencing (scRNA-seq). Amongst tongue innate lymphoid cells, e.g. ILC2, and the specific presence of mast cells in early postnatal tongues, we further identified two main *Irf8*-independent macrophage populations, which were characterized by *Cx3cr1* and *Folr2* expression, respectively. *Cx3cr1*-expressing macrophages were specifically enriched in the lamina propria of the tongue and were detected in fungiform papillae, which harbor taste buds, but were absent from the epidermis. *Folr2*-expressing tongue macrophages localized in muscular tissue and in the lamina propria. These anatomical niches were colonized during embryonic and early postnatal development from a *Cx3cr1*-expressing precursor of high proliferation capacity. Both macrophage populations showed a robust inflammatory response after *in vivo* lipopolysaccharide (LPS) administration, including shared and unique pathways.

In summary, our data provide a detailed atlas of the immune cells of the tongue that will facilitate future research of this under-investigated barrier organ.

## Results

### Characterization of murine tongue hematopoietic cells

To examine the tissue-resident immune compartment of the tongue in an unbiased manner, we performed scRNA-seq of FACS-purified CD45^+^ hematopoietic tongue cells isolated from PBS-perfused adult wild-type C57BL/6 mice. Two biologically and technically independent 10X Chromium experiments were performed that yielded highly reproducible results (**Suppl. Fig. 1a+b**).

**Figure 1:**
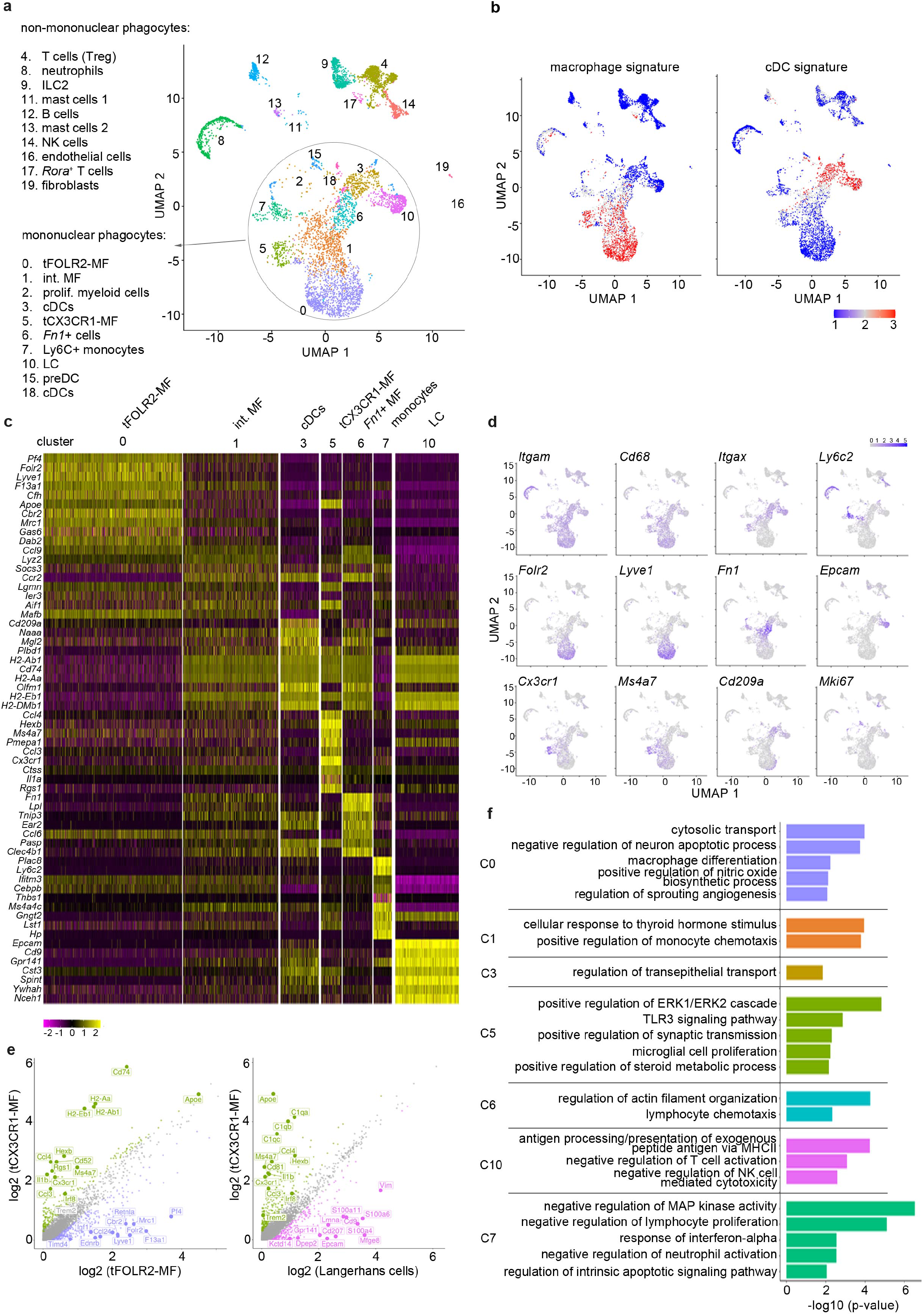
Characterization of mouse tongue leukocytes (**a**) UMAP representation of 6773 sequenced tongue leukocytes from adult, female Bl6 mice (pool of n=8 mice). Data from a biologically and technically independent experiment are shown in Suppl. Fig. 1a+b. Cluster annotation was performed with SingleR (Suppl. Fig. 1c). See Suppl. Data 1+2 for complete gene lists and marker genes for all clusters. (**b**) Gene Set Variation Analysis (GSVA) analysis for the discrimination of macrophages and dendritic cells. One signature gene list for macrophages (derived from ^16^) and one for cDC (derived from ^17^) were used to evaluate the enrichment score for each list in the identified 19 clusters. See also Suppl. Fig. 1d for cDC1 and cDC2 gene signatures and Suppl. Data 3 for full gene lists. Cells with the highest similarity to the signature are labeled red. (**c**) Heatmap of top marker genes for the main mononuclear phagocyte clusters. See Suppl. Fig. 1b for a heatmap of marker genes for all clusters (**d**) Expression pattern of example genes laid over the UMAP for dimension reduction from Figure 1a. (**e**) Differentially expressed genes in tFOLR2-MF vs. tCX3CR1-MF (left) and tCX3CR1-MF vs. tongue Langerhans cells (right). Indicated genes show an increased expression of > 1.5 with an adjusted p-value < 0.05. (**f**) Gene ontology analysis of the differential expressed genes. Redundant pathways were excluded from representation. Only GO annotations involved in biological processes are shown. See Suppl. Data 4 for full list of GO terms per cluster.

We sequenced a total of 6773 cells that clustered into 19 transcriptionally distinct subsets (**Fig. 1a+b**) and used singleR ^14^ for cell lineage recognition (**Suppl. Fig. 1c**). A full list of marker genes and average expression values per cluster can be found in **Suppl. Data 1+2**. We detected type 2 innate lymphoid cells (cluster 9: ILC2 defined by *Gata3*, *Rora*, *Ctla2* and *Arg1*), regulatory T cells (cluster 4; characterized by *Ikzf2*, *Tnfrsf18*, *Cd3d* and *Cd28*), *Rora^+^* T cells (cluster 17; characterized by *Cd3e*, *Lck*, *Cd28* and *Il2rb*), natural killer cells (cluster 14; defined by *Cd3d*, *Nkg7*, *Klrd1*, *Xcl1* expression), B cells (cluster 12; characterized by *Cd79a* and *Cd19*), mast cells (clusters 11 and 13; defined by *Kit* and *Ms4a2*), neutrophils (cluster 8; defined by *S100a9* and *Retnlg*), a few endothelial cells (cluster 16, which is mainly present in subsequent scRNA-seq analysis of Fig.3; defined by *Aqp1*, *Col4a1* and *Pecam1*), fibroblasts (cluster 19; defined by *Dcn*, *Peg3*, *Cald1*), and small clusters of proliferating precursor cells (*Mki67*, *Mcm* gene family and *Top2a*; clusters 7, 14 and 16). However, the majority of cells (65%) fell into the broad category of mononuclear phagocytes and included Langerhans cells (defined by *Epcam* and *Cd9* expression; cluster 10), different classical dendritic cell (cDC) subsets (cluster 3, 15, 18 and 6), monocytes (characterized by *Ly6c2*, *Cebpb*, *Nr4a1* expression; cluster 6) and three subsets of *Cd68-* expressing macrophages (clusters 0, 1 and 5).

As mononuclear phagocytes would be the first responders in tongue immunity, we focused our subset analysis on this major cell compartment. First, we separated mononuclear phagocytes into macrophages and dendritic cells (DC). For this, we performed a gene set variation analysis (GSVA) ^15^ by taking advantage of published cDC and macrophage gene signatures ^16,17^ (**Suppl. Data 3**). The transcriptional signature of clusters 3, 15 and 18 correlated with the cDC gene signature (**Fig. 1b**) and they could further be separated into cDC1 (cluster 3; defined by *Cd209a*, *Cd24a* and *Irf8*) and DC precursors (clusters 15 and 18; characterized by *Hmgb2*, *Asf1b*, *Atad2*). While singleR annotated the *Fn1*, *Lpl*, *Ear2* expressing cells with MHCII-related gene expression in cluster 6 as CD11b^+^ DCs (**Suppl. Fig. 1c**), these cells had neither a strong DC nor a strong macrophage signature (**Fig. 1b**). As the specific cDC2 gene signature ^17^ was also absent in this cluster (**Suppl. Fig. 1d**), the precise ontogeny of these cells will need further investigation.

The GSVA analysis revealed three clusters of cells with macrophage identity, of which clusters 0 and 5 are likely end-stage differentiated macrophage subsets (**Fig. 1b**). Cells in the third macrophage cluster (cluster 1) seem to be macrophages of intermediate differentiation, as their transcriptomic signature shares features with both cluster 0 and 5 (**Fig. 1c+d**). Of the two terminally differentiated macrophage clusters, cluster 0 (hereafter referred to as tFOLR2-MF) expressed high levels of *Folr2*, *Lyve1*, *Pf4* and *Timd4*, while cluster 5 (hereafter referred to as tCX3CR1-MF) expressed high levels of *Cx3cr1*, *Hexb*, *Ms4a7*, *Itgax* and *Pmepa1* (**Fig. 1c+d**). A pairwise comparison of tFOLR2-MF and tCX3CR1-MF transcriptomes revealed 602 differentially expressed genes (DEGs), indicating major differences between the two tongue macrophage populations (abs(FC) > 1.5 and adjusted p-value < 0.05) (**Fig. 1e**). Both tCX3CR1-MF and tFOLR2-MF were transcriptionally distinct from tongue Langerhans cells (tLCs; **Fig. 1e**), which rather fell under the cDC signature (**Fig. 1b**)

We performed gene ontology (GO) enrichment analysis for the different mononuclear phagocyte clusters to gather information on possible distinct functions of these transcriptionally defined cell subtypes. Marker genes of intermediate clusters 1, 3 and 6 were not particularly enriched for any GO biological processes, which potentially reflects their intermediate gene expression signature (**Fig. 1e**; the full list of GO annotation is listed in **Suppl. Data 4**). tFOLR2-MF on the other hand were enriched for gene sets associated with blood vessel biology (“regulation of sprouting angiogenesis”) and macrophage function (“cytosolic transport”, “macrophage differentiation” and “positive regulation of nitric oxide biosynthesis”); tCX3CR1-MF showed gene enrichment for broad immune biological processes such as “positive regulation of ERK1/2 cascade” or “TLR3 signalling pathway” and exhibited similarities to CNS-resident microglia with gene enrichment in the biological process “microglial cell proliferation” (**Fig. 1f**).

Recent studies have indicated potential interactions of *Cx3cr1*-expressing macrophages with neurons ^18–20^. However, unlike these CX_3_CR1^+^ brown adipose tissue and skin nerve-associated macrophages ^18–20^, we did not detect a specific and significant enrichment for genes involved in axon guidance, such as *Plexina4* in tCX3CR1-MF (**Suppl. Fig. 1e**).

Altogether, our scRNA-seq data show that the tongue harbours a wide range of tissue-resident immune cells, of which the majority belong to the mononuclear phagocyte system. We identified two terminally differentiated macrophage subsets: tCX3CR1-MF that have a transcriptomic signature associated with innate immune signaling and tFOLR2-MF that seem to function in blood vessel biology and phagocytosis.

### Tongue macrophages belong to the family of interstitial macrophages

We next established a protocol for the identification and isolation of tCX3CR1-MF and tFOLR2-MF by flow cytometry in *Cx3cr1*^Gfp/+^ reporter mice ^21^. After DNase/Collagenase IV/Hyaluronidase digestion of the tongue (**Suppl. Fig. 2a**), we were able to detect CD64^+^ cells that could further be separated into cells expressing high levels of *Cx3cr1*-GFP (tCX3CR1-MF) and cells that stained positive for Folr2 (tFOLR2-MF; **Fig. 2a**). tCX3CR1-MF also expressed the surface receptors CX_3_CR1, F4/80, MHCII and CD11c, while tFOLR2-MF were additionally characterized by LYVE1 and TIMD4 expression (**Fig. 2b**). These surface characteristics could also be used to identify the tCX3CR1-MF and tFOLR2-MF in WT Bl6 animals.

**Figure 2:**
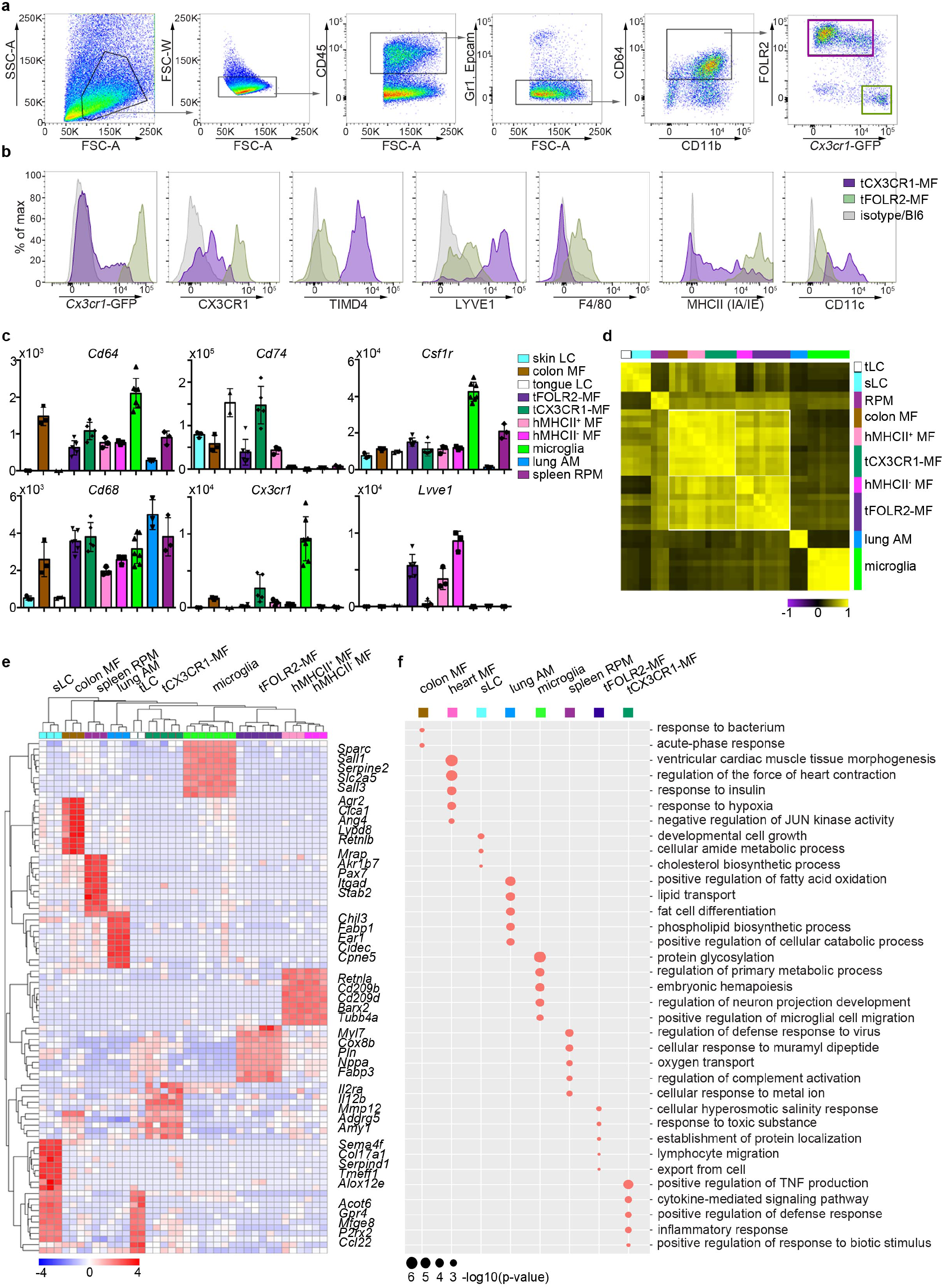
Tissue-specific transcriptomic identity of tongue macrophages Exemplary flow cytometry analysis for tCX3CR1-MF and tFOLR2-MF in *Cx3cr1*^Gfp/+^ mice. Shown are histogram expression patterns for *Cx3cr1*-Gfp, CX3CR1, TIMD4, LYVE1, F4/80, MHCII and CD11c on tFOLR2-MF (violet) and tCX3CR1-MF (green). Either isotype controls or wild-type Bl6 mice were used to control for antibody stain or GFP signals, respectively. (**c**) Different tissue resident macrophage populations were isolated by FACS (see Suppl. Fig. 3 for gating strategy) and analyzed by bulk RNA sequencing. Normalized read counts for important macrophage genes are shown across subsets. See also Suppl. Data 5 for normalized read counts. (**d**) Sample-wise expression correlation analysis of the different macrophage subsets is shown. Color code as indicated in c. (**e**) Heatmap of upregulated genes across macrophage populations. The top 10 upregulated genes per population compared to all other populations are depicted. See also Suppl. Data 6 for full list of upregulated genes. (**f**) Shown are GO annotations of biological processes that are enriched in specific macrophage subsets. Note that tLC showed no specific enrichment and are therefore not represented in the graph.

To identify tongue-specific signatures of tCX3CR1-MF and tFOLR2-MF and to place them in the context of macrophage biology, we compared the transcriptional profiles of FACS-purified tongue macrophages with macrophages isolated from other tissues. We FACS-isolated microglia from the brain, alveolar macrophages from the lung (lung AM), heart MHCII^+^ and MHCII^−^ *Cx3cr1*-Gfp^+^ macrophages, MHCII^+^ intestinal macrophages (colon MF) and F4/80^+^ splenic red pulp macrophages (spleen RPM; see **Suppl. Fig. 3** for gating strategy and **Suppl. Data 5** for full read count table). We additionally sorted skin and tongue Langerhans cells for comparison (sLC and tLC, respectively). Of note, a different digestion protocol was necessary to isolate Epcam^+^ tLCs from the epithelial layer of the tongue (**Suppl. Fig. 2b**).

**Figure 3:**
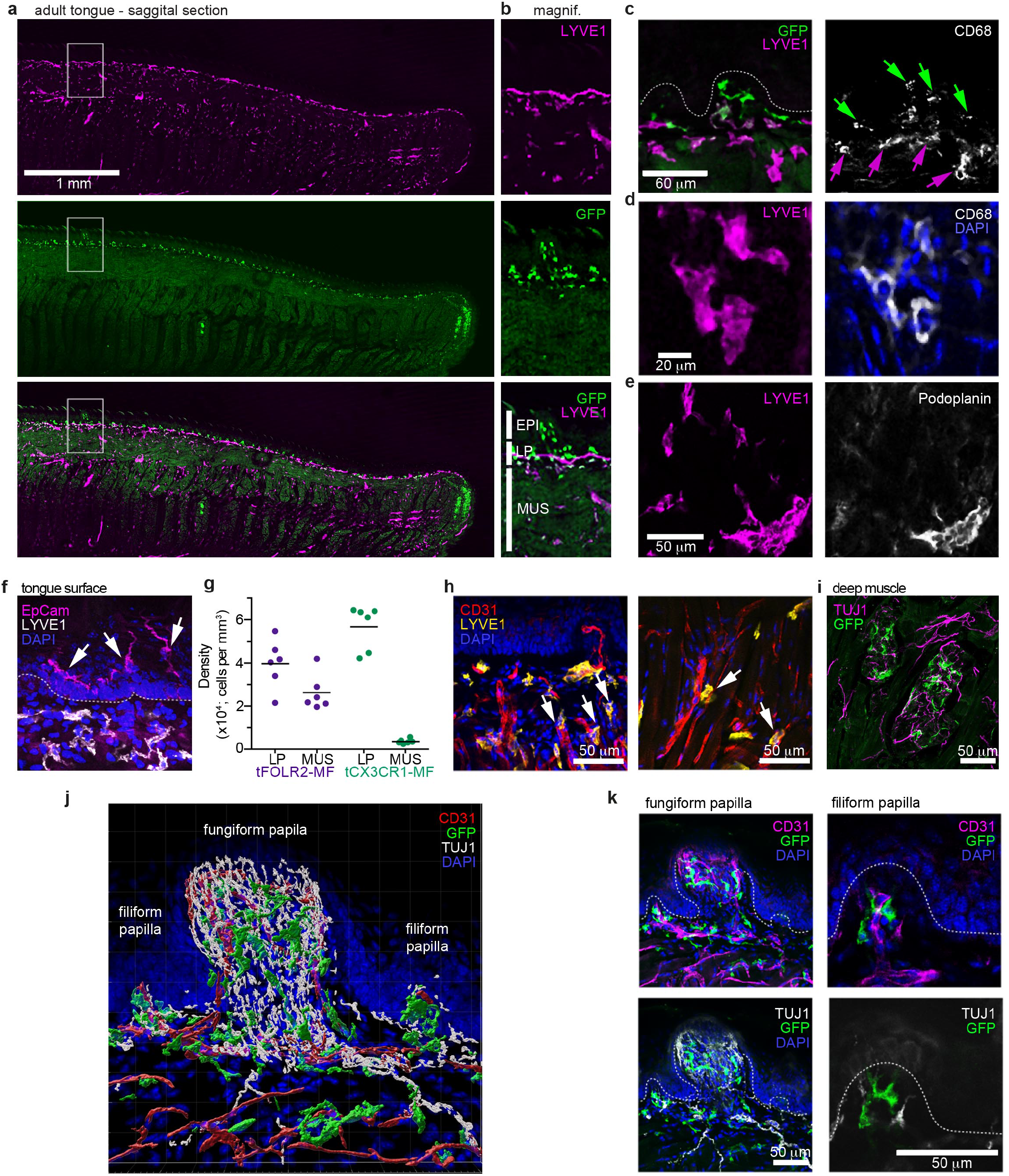
Distinct localization of mouse tongue macrophages (**a**) Panoramic image of a tongue sagittal section from a perfused *Cx3cr1*^Gfp/+^ mouse. The tongue was stained for anti-GFP (CX3CR1; green) and anti-LYVE1 (violet). (**b**) Magnification of the area depicted in (a) that includes a fungiform papilla with annotation of different tissue layers. EPI = epidermis, LP = lamina propria, MUS = muscle. See also Suppl. Fig. 4 for definition of layers. (**c**) Co-staining of anti-GFP (green) and anti-LYVE1 (violet) with anti-CD68 (white). The dashed white line indicates the border between the epidermis and the lamina propria. Green arrows indicate *Cx3cr1*-GFP^+^CD68^+^ cells, while violet arrows highlight LYVE1^+^CD68^+^ macrophages. (**d**) High magnification of LYVE1^+^ cells (violet) stained with anti-CD68 (white) and DAPI (blue). (**e**) Discrimination of podoplanin^+^ (white) lymphatics from LYVE1^+^ (violet) podoplanin^−^ cells. (**f**) Anti-Epcam (violet) staining identifies Langerhans cells in the epidermis of the tongue. Sections were stained with anti-Lyve1 (white) and DAPI (blue). (**g**) Quantification of tFOLR2-MF and tCX3CR1-MF in lamina propria (LP) and muscle layer (MUS) in adult female mouse tongues. Each dot represents one animal. See also Suppl. Fig. 4. (**h**) Proximity of LYVE1^+^ (yellow) cells to CD31^+^ (red) blood vessels. (**i**) Rare *Cx3cr1*-GFP^+^ (green) cell clusters could also be detected in innervated Tuj1^+^ (violet) areas in posterior regions of the tongue. (**j**) 3D reconstruction of *Cx3cr1*^Gfp/+^ tongue tissue sections stained for anti-CD31 (red), anti-Tuj1 (white), anti-GFP (Cx3cr1; green) and DAPI (blue). (**k**) Localization of tCX3CR1-MF in fungiform (right) and at the base of filiform (left) papillae. Sections were stained for anti-CD31 (violet), anti-Tuj1 (white), anti-GFP (Cx3cr1; green) and DAPI (blue).

To validate the data quality, we compared expression of common macrophage-related genes in the different macrophage populations (**Fig. 2c**). All cells except Langerhans cells expressed the macrophage genes *Cd68*, *Mertk* and *Fcgr1*. Furthermore, *Csf1r* expression was detected in almost all macrophage subsets with particularly high levels in microglia, but not in alveolar macrophages ^16^. *Cx3cr1*, *Lyve1* or MHCII-related genes such as *Cd74* were also expressed according to the expected expression pattern (**Fig. 2c**), which confirmed the accuracy of our gating and sorting strategy.

Correlation analysis of all macrophage populations revealed that, in line with published data ^14,22^, classical TRM populations such as microglia, splenic macrophages, Langerhans cells and alveolar macrophages each had a very distinct expression profile, indicating the robust tissue imprinting of these cells (**Fig. 2d**). Tongue macrophages on the other hand were more related to heart and intestinal macrophages, regardless of their tissue of residence. Morover, even within this group of TRM, MHCII expressing cells like heart MHCII^+^ macrophages, tCX3CR1-MF and colon macrophages showed a higher correlation to each other and were distinct from MHCII^−^ cell populations, including tFOLR2-MF and heart MHCII^+^ macrophages (**Fig. 2d**). Thus, tCX3CR1-MF and tFOLR2-MF share similarities with the two main interstitial macrophage populations that have been previously identified across various tissues ^19,22^

We focused our analysis on up-regulated genes (FC > 2; adjusted p-value < 0,001) to identify a tissue-specific signature of each macrophage subset and were able to annotate previously described marker genes to macrophage populations isolated from the lung, heart, skin, brain and spleen ^16,23^. tCX3CR1-MF on the other hand expressed significantly higher levels of *Il2ra*, *Il1a* and *Irak2* compared to all other tested TRM subsets, while tFOLR2-MF were characterized by the transcription of *Cd209* gene family members (*Cd209b*/*d*/*f*), *Retnla*, *Clec10a* and *Fxyd2*. The top genes for each macrophage subset are shown as a heatmap in **Fig. 2e** and a full list of these up-regulated genes can be found in **Suppl. Data 6**.

We next performed a GO enrichment analysis on these DEGs and found in agreement with previously published work ^24^ an enrichment of GO terms that facilitate the tissue-specific function of each TRM subset (**Fig. 2f**). In comparison, tCX3CR1-MF showed a strong enrichment for genes involved in inflammatory pathways, such ’positive regulation of TNF production’ or ’cytokine-mediated signaling pathway’, which is in line with our scRNA-seq data presented in **Fig. 1**. tFOLR2-MF were characterized by weak but significant enrichment for ’cellular hyperosmotic salinity response’ and ’response to toxic substances’.

Taken together, these data demonstrate that tCX3CR1-MF and tFOLR2-MF tongue macrophages belong to the family of interstitial macrophages. They fall into the two broad categories of *Cx3cr1*- and *Lyve1*/*Folr2*/*Timd4*-expressing cells ^19,22^, but also show unique transcriptomic signatures that probably reflect the requirements of their local tissue niche.

### Distinct localizations of tCX3CR1- and tFOLR2-macrophages in the adult tongue

The unique transcriptomic signatures of tCX3CR1-MF and tFOLR2-MF could indicate that these populations inhabit different microanatomical niches within the tongue. To test this hypothesis, we performed immunohistochemistry on adult mouse tongues. *Cx3cr1*^Gfp/+^ animals were perfused and fixed tongue sections were stained with antibodies against GFP and LYVE1. Of note, we have used LYVE1 and FOLR2 markers interchangeably for the identification of tFOLR2-MF. Indeed, GFP^+^ cells were concentrated in the lamina propria, while LYVE1 staining was evident on cells throughout the tongue tissue with the exception of the epidermis (**Fig. 3a+b**). Since lymphatic vessels also stain positive for LYVE1 ^25^, the tissue was counterstained with antibody against CD68, a common pan-macrophage marker. Thus, we could identify tCX3CR1-MF as double-positive CD68^+^*Cx3cr1*-GFP^+^ cells in the lamina propria, but not in the tongue epidermis or muscle (**Fig. 3c**) and tFOLR2-MF as double-positive CD68^+^LYVE1^+^ cells in the lamina propria and the underlying muscle (**Fig. 3d**). Of note, tFOLR2-MF were morphologically distinct from LYVE1^+^Podoplanin^+^ lymphatics (**Fig. 3e**). EPCAM^+^ Langerhans cells with a ramified morphology localized exclusively in the epidermis of the tongue (**Fig. 3f**).

To better characterize the tissue localisation of tCX3CR1-MF and tFOLR2-MF, we quantified their distribution in different layers of the tongue. CD68^+^LYVE1^+^ double-positive tFOLR2-MF localized in the muscular layer as well as in the lamina propria (**Fig. 3g+h** and **Suppl. Fig. 4**), while CD68^+^*Cx3cr1*-GFP^+^ double-positive tCX3CR1-MF were only detected in the lamina propria and were virtually absent in muscular tissue (**Fig. 3g**). However, clusters of *Cx3cr1*-GFP^+^ cells could be detected in the posterior part of the tongue, along Tuj1^+^ nerves that possibly cater to circumvallate and foliate papillae (**Fig. 3i**) and in innervated areas of the deep muscle, along the chorda tympani branch of the facial nerve. tCX3CR1-MF were present at the base of both filiform and fungiform papillae (which harbour taste buds) and within the lamina propria, which is densely innervated by sensory fibres (**Fig. 3j+k**).

**Figure 4:**
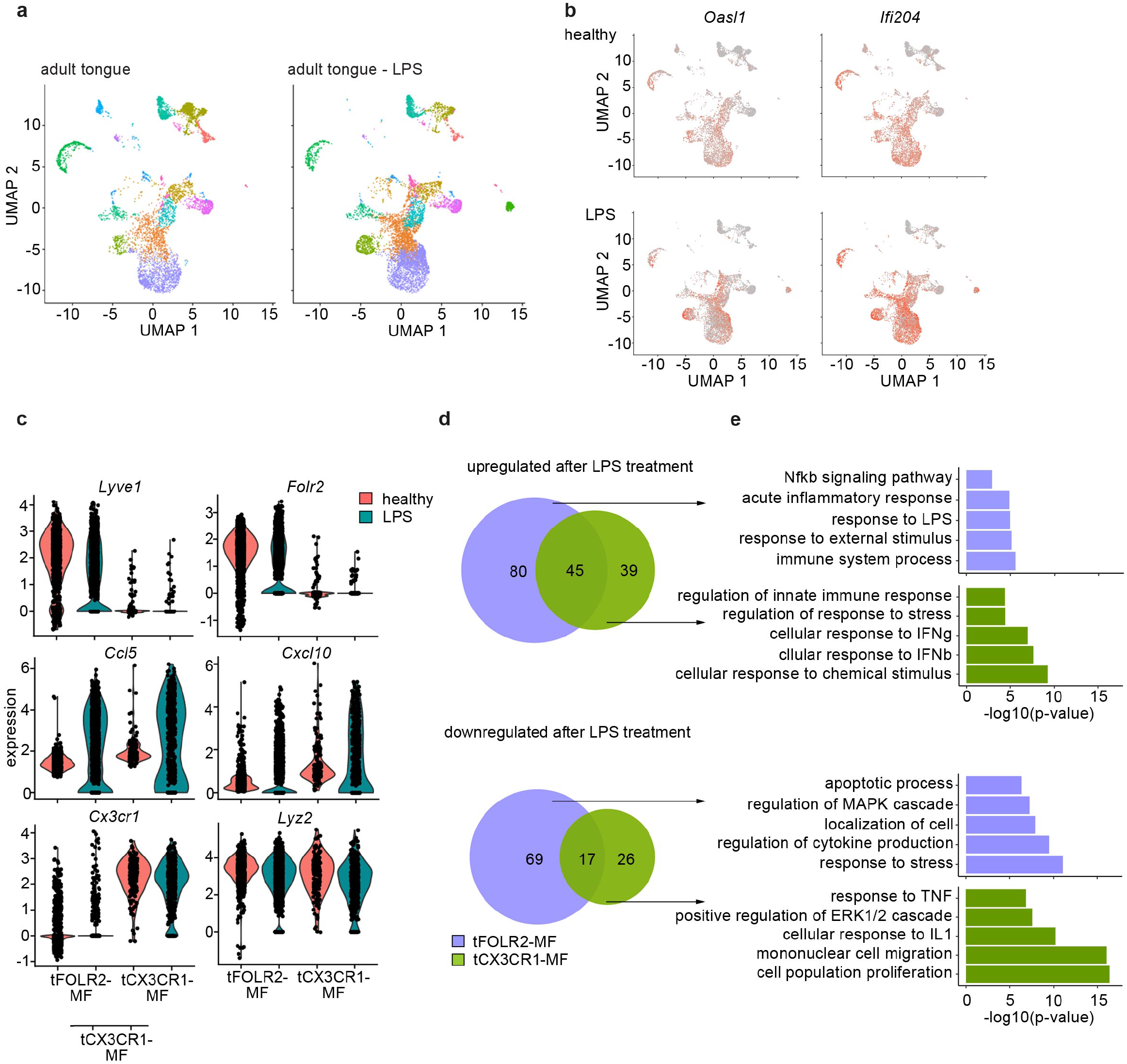
Inflammatory response of tongue macrophages to systemic LPS challenge (**da**) Female Bl6 mice were intraperitonally injected with 1 mg/kg LPS and tongue CD45^+^ leukocytes were FACS purified 6 hours after injection (pool of 6 mice). In total, 8165 LPS-exposed tongue leukocytes were sequenced and the data was integrated with the existing adult analysis in order to allow population comparison. Shown are UMAPs for adult cells (left) and LPS-treated cells (right). (**b**) Gene expression pattern of *Oasl1* and *Ifi204* in healthy and LPS-treated tongue leukocytes. (**c**) Violin blots showing marker gene expression patterns in untreated (red) and LPS treated (green) tFOLR2-MF and tCX3CR1-MF. (**d**) Number of up- and down-regulated genes in tFOLR2-MF (violet) and tCX3CR1-MF (green) after LPS injection. The full list of DEGs can be found in Suppl. Data 7. (**e**) GO enrichment analysis of the DEGs. Only GO annotations involved in biological processes are shown.

Thus, we show here that tongue tCX3CR1-MF and tFOLR2-MF inhabit distinct anatomical regions of the tongue. tCX3CR1-MF localized in the highly innervated lamina propria at the base of filiform papillae and within fungiform papillae, while tFOLR2-MF can additionally be found in deeper layers, often in proximity to blood vessels.

### Response of tongue macrophages to systemic inflammation

Regardless of their tissue-specific roles in homeostasis, macrophages usually also function as first-line responders to pathogens. We therefore tested the inflammatory response of tongue immune cells to the bacteria endotoxin lipopolysaccharide (LPS). Mice were challenged intraperitoneally (i.p.) with LPS and transcriptomic changes of the tongue hematopoietic system (CD45^+^ cells) were determined by scRNA-seq 6h later. In total, we sequenced 8165 cells. Integration of the LPS data with the data from steady state mice indicated that all cell populations were present 6 hours after LPS injection (**Fig. 4a**). LPS injection led to a general increase of inflammatory gene expression such as *Oasl1* and *Ifi204* across all cells of the tongue immune system (**Fig. 4b**).

We focused on tongue-resident macrophages of cluster 0 (tFOLR2-MF) and cluster 5 (tCX3CR1-MF) and examined DEGs between the steady state and LPS conditions in these two populations. 211 DEGs were identified in tFOLR2-MF (the full list of DEGs can be found in **Suppl. Data 7**), of which 125 genes were upregulated (e.g. *Il1b*, *Relb* and *Slfn4*) and 86 genes downregulated after LPS injection (e.g. *Folr2*, *Lyve1* and *Klf4*; **Fig. 4c+d**). Interestingly, many of the tFOLR2-MF signature genes (i.e. *Folr2* and *Lyve1*) were downregulated after LPS exposure (**Fig. 4c**). In tCX3CR1-MF we detected 127 DEGs between the physiological and the pathological state of which 84 were upregulated (e.g. *Ifit2*, *Lgals3* and *Usp18*) and 43 were downregulated (e.g. *Lyz2*, *Ccr2* and *Cd9*). LPS induced upregulation of 45 common genes (e.g. *Cxcl10*, Ccl5, *Il1rn* and *Gbp2*) and downregulation of 17 common genes in both tongue macrophage subsets (e.g. *Fcrls*, *S100a10*, *Lyz1* and *Retnla*; **Fig. 4c+d**).

GO enrichment analysis was used to explore potential signaling differences of tFOLR2-MF and tCX3CR1-MF in response to systemic LPS. Shared upregulated genes in the two subsets were involved in ‘defense response’, ‘response to cytokine’ and ‘response to bacterium’. Genes that were only upregulated in tCX3CR1-MF macrophages were particularly enriched for GO terms associated with type I interferon signaling (**Fig. 4e**). On the other hand, tCX3CR1-MF showed downregulation of genes involved in ‘mononuclear cell migration’ and ‘cell population proliferation’ after LPS exposure (**Fig. 4e**). tFOLR2-MF showed the specific upregulation of genes involved in ‚immune system process’ and ‚Nfkb signaling’, while they downregulated in response to LPS the ‘response to stress’, ‚localisation of cell’ apoptotic processes.

Thus, both tFOLR2-MF and tCX3CR1-MF were activated by systemic administration of bacterial components (LPS). However, they responded differently, with strong type I interferon signaling response characterizing tCX_3_CR1-MF and a more ‘classical’ Nfkb-mediated macrophage response seen in tFOLR2-MF.

### Distribution and subset analysis of tongue macrophages during development

We next investigated the spatiotemporal distribution of tongue macrophages over development in an effort to shed light on the origin and timeline of establishment of tFOLR2-MF and tCX3CR1-MF populations. First, we isolated leukocytes from Cx3cr1^Gfp/+^ reporter mice at different ages and performed flow cytometry to detect CD11b^+^CD64^+^ macrophages that we could further separate according to *Cx3cr1*-GFP expression and FOLR2 immunoreactivity. We used FOLR2 as a marker since macrophages at this developmental stage do not show LYVE1 surface expression. At embryonic day 17.5 (E17.5), all tongue macrophages were characterized by high *Cx3cr1*-GFP expression (**Fig. 5a**). Of these *Cx3cr1*-GFP^+^ cells, two-thirds additionally expressed FOLR2 (G2 in **Fig. 5a**). Similar proportions of macrophage subsets were observed in the mouse tongue at postnatal day 3 (p3). At this time-point, an intermediate *Cx3cr1*-GFP^int^ FOLR2^+^ subset was also present (G3 in **Fig. 5a+b**). In subsequent developmental stages (p11 and p28), the proportion of double-positive *Cx3cr1*-GFP^+^FOLR2^+^ cells progressively decreased and two main *Cx3cr1*-GFP^+^FOLR2^−^ and *Cx3cr1*-GFP^int^FOLR2^+^ macrophage populations (corresponding to tCX3CR1-MF and tFOLR2-MF respectively) were established by 8 weeks of age (**Fig. 5b**).

**Figure 5:**
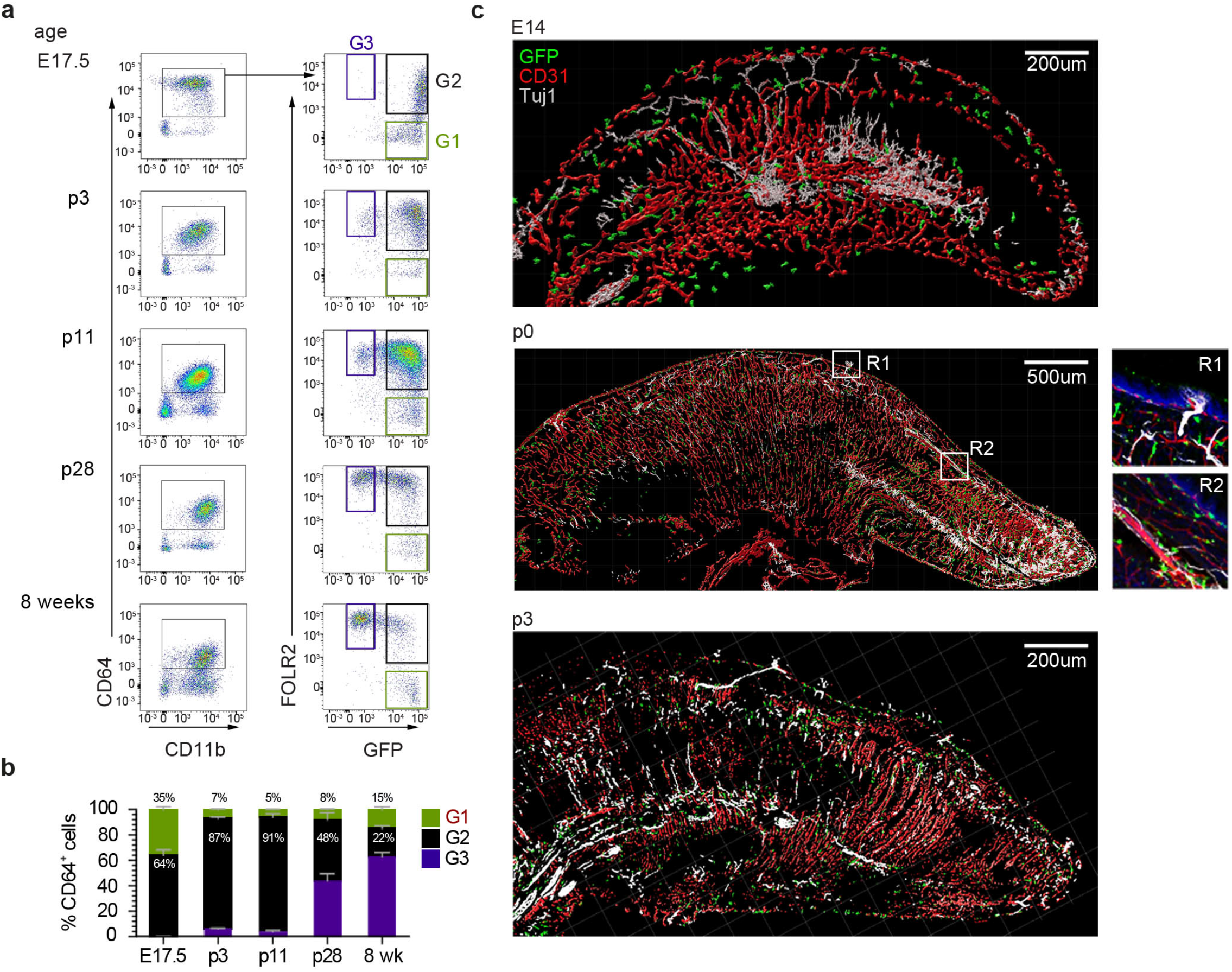
Tongue macrophage distribution during development (**a**) Flow cytometry analysis of tongue leukocytes isolated from *Cx3cr1*^Gfp/+^ mice at different embryonic (E) and postnatal (p) stages of development. Cells were pre-gated for CD45^+^ Epcam^−^ Gr1^−^. (**b**) Distribution of *Cx3cr1*-GFP^+^ FOLR2^−^ (G1), *Cx3cr1*-GFP^+^ FOLR2^+^ (G2) and *Cx3cr1*-GFP^−^ FOLR2^+^ (G3) cells out of CD64^+^ cells. ^5–7^ animals per time point were used. (**c**) 3D reconstructions of sagittal tongue sections stained with anti-GFP (green), anti-CD31 (blood vessels; red) and anti-TUJ1 (neurons; white) from E14, p0 and p3 Cx3cr1^Gfp/+^ reporter mouse tongues.

To correlate the flow cytometry data to spatial localization, we performed immunohistochemical analysis of tongue sections from *Cx3cr1*^Gfp/+^ mice during early developmental stages. As mentioned above, tongue macrophages did not express *Lyve1* during development and the FOLR2 antibody used for FACS did not work for immunohistochemical staining of the tissue. We thus relied on *Cx3cr1*-GFP as a marker of macrophages. Tissue sections were counterstained with antibodies against CD31 to identify blood vessels, and, since various nerves (e.g. VII^th^, IX^th^ and X^th^ cranial nerve ganglia) innervate the tongue tissue during embryogenesis, we also stained tissue sections for anti-beta Tubulin III (Tuj1) to visualize neurons. At E14.5 *Cx3cr1*-GFP^+^ macrophages could be detected throughout the whole tongue tissue with no specific localization pattern (**Fig. 5c**). This continued through postnatal day p0 and until p10, whereby *Cx3cr1*-GFP^+^ macrophages were still dispersed throughout the tongue but started to align along the lamina propria. At these stages, taste bud maturation is observed ^26,27^ and we detected the first *Cx3cr1*-GFP^+^ macrophages in proximity to fungiform papillae (**Fig. 5c**).

These data demonstrate the progression of a dynamic tongue macrophage compartment during mouse tongue development and that the specific localization of tongue macrophages in distinct anatomical niches is only established during postnatal stages.

### Molecular characterization of tongue macrophages during development

The histological analysis reveals the unordered distribution of *Cx3cr1*-GFP^+^ macrophages during early developmental stages towards distinct localization at adulthood. However, it is not clear from our histological data whether this reflects a sequential developmental maturation of macrophages or rather a progressive exchange of embryonic populations with adult, bone marrow-derived monocytes.

To gain further insight into this question, we investigated the tongue immune compartment from mice at post-natal day 3 (p3) with scRNA-seq. We chose p3 since it is the stage at which we could first identify *Cx3cr1*-GFP^int^FOLR2^+^ macrophages by flow cytometry (**Fig. 5b**). We sequenced 13898 cells (from a pool of n=7 mice) and overlaid the data on the scRNA-seq data derived from adult mice (**Fig. 6a**). All adult tongue immune cell populations were also present at p3, however, we noticed major differences in the frequency of several populations. Notably, p3 tongues harbored more mast cells (clusters 11 & 13) and fewer *Fn1*+ myeloid cells (cluster 6) than adult tongues (**Fig. 6b**). We also detected a newly appearing cluster of cells in p3 tongues (cluster 2), which was almost absent in adult mice (**Fig. 6b**). These cluster 2 cells expressed high levels of proliferation-related genes, including *Top2a*, *Ccnb2*, *Mki67*, and further showed expression of typical macrophage lineage genes such as *Cx3cr1*, *Tlr4*, *Pf4* and *Lyve1*, which suggests they represent proliferating macrophage precursor cells (**Fig. 6c**). Indeed, when the UMAP is projected in 3D, cluster 2 cells seemed to incorporate into tFOLR2-MF (**Fig. 6d; Suppl. Data 8**). To investigate the connection of cluster 2 precursors with tongue macrophages we used Slingshot, which models developmental trajectories in scRNA-seq data ^28^. Slingshot analysis revealed a possible bifurcation of the precursor cells at the cluster 1 level at which trajectories either split into cluster 0 or cluster 5 cells (**Fig. 6e**), which might indicate that both tFOLR2-MF and tCX3CR1-MF populations derive from a common precursor.

**Figure 6:**
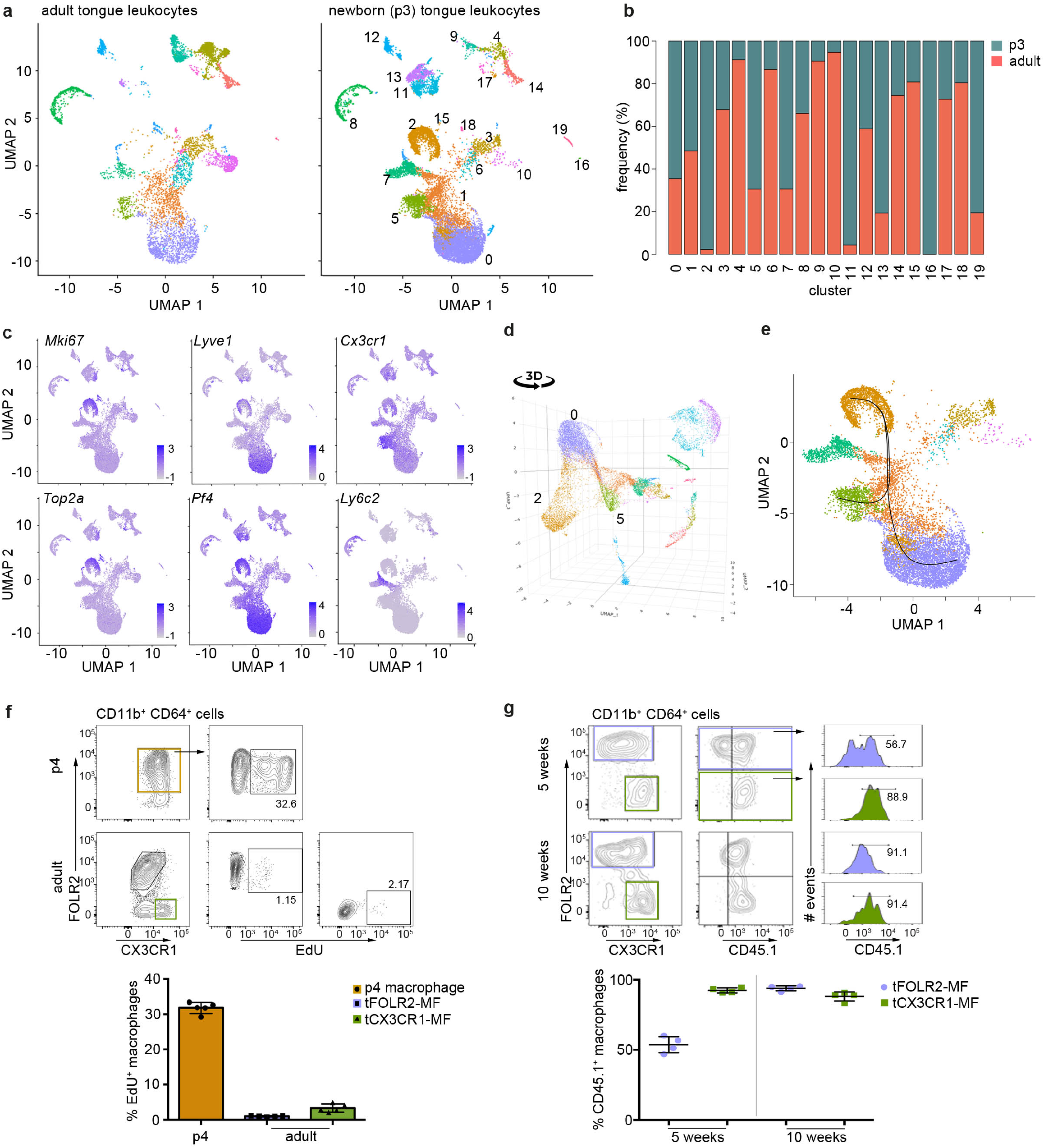
Transcriptomic analysis of hematopoietic tongue cells from newborn mice (**a**) CD45^+^ tongue leukocytes were isolated by FACS from newborn p3 mice. The purified cells were subjected to scRNA-seq and a total of 13898 cells were sequenced. The early postnatal data was integrated with the adult scRNA-seq dataset shown in Fig. 1 for population comparison. Shown are UMAPs for adult (left) and newborn cells (right). (**b**) Cell frequencies for each cell cluster in adult and newborn mice. (**c**) Expression levels of example genes in newborn cells shown in the UMAP. (**d**) 3D UMAP visualization reveals the merging of proliferating cluster 2 cells with cluster 0 cells. The 3D html file is provided in Suppl. Data 8. (**e**) Slingshot analysis was used to model the developmental trajectories of tongue macrophages. (**f**) *In vivo* EdU proliferation assay of adult and newborn tongue leukocytes. EdU was injected and tongue cells were isolated 24 hours after injection. Each dot represents one independent animal. The experiment was performed once. (**g**) CD45.2/2 animals were lethally irradiated (9.5 Gy) and reconstituted with CD45.1/1 bone marrow cells. 5 weeks (left) and 10 weeks (right) after transfer, tongue cells were isolated and investigated by flow cytometry for the distribution of CD45.1+ cells within the macrophage subsets. Each dot represents one independent animal. The experiment was performed once.

We confirmed the proliferation activity of macrophage precursor cells at p3 by EdU *in vivo* labelling. One day after injection, EdU incorporation could be readily observed in about 30-40% of CD68^+^FOLR2^+^ cells, while both adult tongue macrophage subsets showed no signs of homeostatic proliferation (**Fig. 6f**).

We then investigated whether peripheral cells like monocytes could adapt to the tongue macrophage niches in an immune compromised condition, e.g. after whole body irradiation. We performed bone marrow (BM) chimeric experiments, in which CD45.1 BM cells were transferred into irradiated CD45.2 recipients. We analyzed the composition of donor versus host tongue macrophages 5 and 10 weeks after transfer. While tCX3CR1-MF were already replaced by monocyte-derived cells 5 weeks after transfer, the tFOLR2-MF still comprised 53% (+/− 6 SD) donor cells and accordingly, showed a slower replacement (**Fig. 6g**). At 10 weeks after irradiation, both tongue macrophage subsets were exclusively of hematopoietic stem cell (HSC) origin. These data indicate that both tongue macrophage niches can be repopulated by myeloid precursors during adulthood.

To further explore the monocyte-dependency of tCX3CR1-MF and tFOLR2-MF under homeostatic conditions, we profiled the tongue-resident immune compartment of *Irf8*-deficient mice, which lack Ly6C^+^ monocytes and show reduced numbers of Ly6C^−^ monocytes ^18,29^. In total, 9047 CD45^+^ cells were profiled from adult *Irf8*-deficient mice (pool of n=5 mice) by scRNA-seq (**Suppl. Fig. 5**). All tongue-resident hematopoietic cell populations that were present in WT animals could also be identified in *Irf8*^−/−^ mice including dendritic cells, tCX3CR1-MF and tFOLR2-MF. In addition to the unchanged mononuclear phagocyte subset composition in *Irf8*^−/−^ mice, the transcriptome of tCX3CR1-MF and tFOLR2-MF was also largely unaffected by the absence of the transcription factor IRF8 (9 DEGs for *Cx3cr1*^+^ and 9 DEGs for *Folr2*^+^ macrophages; **Suppl. Fig. 5**).

Taken together, these experiments indicated that macrophage precursor cells proliferate locally and give rise to both tongue macrophage subsets. However, during adulthood and under immune compromised conditions, tCX3CR1-MF and tFOLR2-MF can be replaced by HSC-derived cells.

## Discussion

Despite the significance of the tongue as a site of interaction between microbes and the host, its cellular immune composition is not well investigated, especially not on the transcriptomic level. In light of the emerging role of different immune cells such as ILCs, regulatory T cells or macrophages in tissue remodeling, nerve surveillance and homeostatic tissue organization ^18,20,30,31^, we characterized the mouse tongue-resident immune cell compartment under physiological conditions. We confirmed the existence of tLCs, cDCs and Ly6C^+^ monocytes in the tongue ^9,32,33^ and additionally identified two new subsets of tongue-resident macrophages, one of which was characterized by *Folr2* and *Lyve1* expression, while the other subset expressed high levels of *Cx3cr1*, MHCII-related genes and *Itgax* (encoding CD11c). The phenotype of these two tongue macrophage populations is reminiscent of the two recently defined interstitial macrophage populations which are present across various organs ^19,22^. In agreement with this, our bulk RNA-seq analysis revealed greater transcriptional similarities of tongue macrophages with interstitial heart macrophages, for example, than with TRMs such as Langerhans cells or microglia.

Chakarov *et al*., further described an association of *Lyve1*^+^ macrophages with blood vessels and *Cx3cr1*^+^ macrophages with nerve fibers ^19^. In the adult tongue, tFOLR2-MF were distributed across both the lamina propria and the deep muscle of the tongue, but were absent from the epithelium. We could also observe the association of tFOLR2-MF to CD31^+^ blood vessels. On the other hand, tCX3CR1-MF are largely restricted to the highly innervated lamina propria and along the fibers of the chorda tympani branch of the facial nerve. In addition, rare clusters of *Cx3cr1*^+^ macrophages could also be detected in innervated posterioir areas of the tongue. It is therefore quite likely that the previously characterized CD11c^+^ “dendritic cells” in the lamina propria of the human and mouse tongues ^7,34^ correspond to the cells described here as tCX3CR1-MF.

The proximity of tCX3CR1-MF to nerves raises the question of whether they perform specific nerve-associated functions. In the skin, large, peripheral nerve-associated *Cx3cr1*^+^ macrophages were recently described ^18^ that were involved in the maintenance of myelin sheath integrity and axon sprouting after injury and had a transcriptional signature related to nervous system functions. This particular signature was absent from tCX3CR1-MF, although tCX3CR1-MF did express some genes that are enriched in microglia (e.g. *Hexb*, *Apoe*) that also reside in close proximity to neurons. Various cranial nerve ganglia innervate the tongue tissue during embryogenesis and axons are guided by multiple chemoattractive factors to the tongue epithelium such as brain derived neurotrophic factor (BDNF) ^35^. It is known that microglia-derived BDNF plays an important role in synapse formation and plasticity in the adult brain ^36^. It is therefore possible that the subsequent postnatal maturation of taste receptor cells and the synaptic interconnectivity with neurons might be influenced by tCX3CR1-MF. The interesting topic of whether and how macrophages interact with nerves in the tongue needs further investigation.

It is also possible that tongue macrophages contribute directly or indirectly to instances of taste dysfunction. Various infections including Covid-19 or middle ear infections ^37–39^, different medical treatment regimens 40 and aging 41 have all been associated with taste disruption. Taste bud maintenance relies on a continuous renewal of differentiated taste receptor cells ^42^ and it has been shown that systemic inflammation like peripheral LPS injection increases TNFα and IL10 production by taste cells ^43,44^, inhibits taste progenitor cell proliferation and interferes with taste cell renewal ^45^. Such a mechanism might explain taste disorders associated with infections in general. We showed that systemic inflammation also causes a direct response from tongue-resident macrophages. We did not investigate the precise contribution of inflammation to taste sensation, yet consider the possibility that tongue macrophages may release cytokines and potentially neurotoxic products that could cause nerve damage. The reverse has also been shown, that tissue macrophages can also mediate neuronal protection and therefore limit neuronal damage upon infection ^46^. Whether and how the various tongue mononuclear phagocytes are involved in taste perception remains to be explored and could be of clinical relevance in conditions such as anorexia or cancer, where appetite- and weight-loss is often aggravated by taste-dysfunctions.

The question remains of the role of tongue immunity in cases where tongue homeostasis is disrupted. It was recently shown that tongue CD163^+^ macrophages infiltrate tumor tissue in squamous cell carcinoma at a high frequency ^5^. CD163^+^ macrophages were interpreted to be “M2 macrophages”, and increased infiltration correlated with worse outcomes compared to patients with a high infiltration of CD11c^+^ “M1” macrophages ^5^. Regardless of whether the “M1/M2” classification really applies to *in vivo* situations ^47^, our data corroborate the notion of the existence of distinct subsets of tongue macrophages, which might respond differently to tumor-specific environmental cues.

We present a comprehensive catalog of immune cells in the murine tongue in physiological conditions and upon LPS-induced systemic inflammation. We identified two novel macrophage subsets in the tongue, namely tCX3CR1-MF and tFOLR2-MF, and place these findings in the context of mammalian macrophage biology. We hope that these data will encourage and support further investigations of the tongue as barrier and as an underrated immunological organ.

## Methods

### Ethics statement, experimental reporting & study design

This study was performed in strict accordance with national and international guidelines for the care and use of laboratory animals (Tierschutzgesetz der Bundesrepublik Deutschland, European directive 2010/63/EU, as well as GV-SOLAS and FELASA guidelines and recommendations for laboratory animal welfare). The animal experiment permission was obtained from the Landesamt für Gesundheit und Soziales (LAGeSo, Berlin). Most of the mice that were used in this study were also part of other experiments, in order to reduce animal experiments and suffering.

### Mice

We used the following mouse strains: C57BL/6N wildytpe mice, B6(Cg)-Irf8tm1.2Hm/J (*Irf8*^−/−^; Jackson laboratory, stock number: 018298), B6.SJL-Ptprca Pepcb/BoyJ (CD45.1/1; Jackson laboratory, stock number: 002014) and B6.129P2(Cg)-Cx3cr1tm1Litt/J (*Cx3cr1*^Gfp^; ^21^). For BM chimeras, 8-12 weeks old recipient animals (CD45.2/2) were lethally irradiated (950 rad) and reconstituted with 10^6^ BM cells isolated from WT (CD45.1/1) BM cells. The animals received Ciproxin in their drinking water for 10 days after irradiation. BM chimeras were analyzed 5 and 10 weeks after transfer. Mice were bred and housed at specific pathogen-free (SPF) animal facilities of the Max-Delbrück-Center for Molecular Medicine in Berlin, Germany or at the Weizmann Institute of Science in Rehovot, Israel. Mice were kept in standard conditions (22±1°C) under a 12-hour light cycle with experiments carried out during the “lights-on” phase. Mice had access to a chow diet *ad libitum* and cages were lined with chip bedding and enriched with a mouse tunnel/igloo.

### Cell suspensions for fluorescence-activated cell sorting

Adult mice were anesthetized by intraperitoneal injection of 150mg/kg body weight pentobarbital sodium (WDT) and intracardially perfused with PBS.

#### Brain

Brains were dissected, and the olfactory bulb and cerebellum were removed. The remaining brain was minced and filtered through a 70μm cell strainer in ice-cold high-glucose DMEM (Sigma). The cell suspension was centrifuged at 1,200 rpm at 4°C for 6 minutes and the pellet was resuspended in 5 ml of 40% Percoll (GE Healthcare) in PBS and transferred to a 15 ml tube. Tubes were centrifuged at 2,000 rpm for 25 minutes (min) at 14°C with no acceleration nor break. Pellet was resuspended in 10 ml sorting buffer and centrifuged for 6 min at 1,200 rpm and 4°C. Leukocyte-containing cell pellets were further processed.

#### Colon

Cells from *Cx3cr1*^Gfp/+^ mice were isolated as previously described ^48^. In brief, the intestine was removed and feces were flushed with cold PBS without calcium and magnesium. The intestine was longitudinally opened and cut into 0,5 cm pieces. Intestinal epithelial cells were removed by incubation with the HBSS containing 1mM DTT, 2mM EDTA and 5% of fetal calf serum. The cell suspension was incubated at 37 °C, 125 rpm for 40 min. The cell suspension was shortly vortexed and passed through a 100μm mesh. The intestinal tissue pieces were transferred to a 50 ml Falcon tube containing 5 ml PBS with 5% FBS, 1 mg/ml of collagenase VIII (Sigma) and 0.1 mg/ml DNase I. Samples were incubated at 37 °C, 250 rpm for 40 min. Digested colon tissue was vortexed for 40 sec, passed through a 80μm mesh and cells were collected by centrifugation at 1,200 rpm for 10 min at 4 °C. Cells were stained after CD16/32 block with antibodies against CD45, CD11b, lineage (Ly6C, Ly6G, B220) and MHCII.

#### Heart

The heart of Cx3cr1^Gfp/+^ mouse was removed, the organ was chopped into small pieces and digested for 30 min at 37°C in RPMI medium without fetal calf serum supplemented with 1 mg /ml Collagenase IV and 1 mg/ml DNase I. The digestion was stopped by addition of staining buffer (2mM EDTA, 1% FCS in PBS) and the cell suspension was minced through a 100 μm cell strainer. After centrifugation at 1,200 rpm, 4°C for 6 min, supernatant was discarded and cells were blocked with anti-CD16/32 antibodies for 10 min, before anti-F4/80-biotin antibodies were added for 25 min on ice. After washing, the cell pellet was incubated with anti-biotin beads (Miltenyi) for 15min on ice. Cells were washed again (1,200 rpm, 4°C for 6 min) and MACS was performed with LS columns (Miltenyi). After elution and washing of cells, lung cells were stained for CD45, streptavidin, dump (Ly6C, Ly6G), CD11b, MerTK, CD64 and MHCII

#### Lung

The lung was removed and cut into small pieces. Tissue was collected in RPMI medium without fetal calf serum supplemented with 1 mg/ml Collagenase A and 1 mg/ml DNase I. Tissue was digested for 30 min at 37°C. The suspension was minced and filtered through a 100μm cell strainer. The cell suspension was centrifuged at 1,200 rpm, 4°C for 6 min. The supernatant was removed and cells were blocked with anti-CD16/32 antibodies for 10 min, before anti-CD45-biotin antibodies were added for 25 min on ice. Afterwards, cells were washed with staining buffer (2mM EDTA, 1% FCS in PBS) at 1,200 rpm, 4°C for 6 min, supernatant was discarded and anti-biotin beads (Miltenyi) were added for 15min. Cells were washed again (1,200 rpm, 4°C for 6 min) and MACS was performed with LS columns (Miltenyi). After elution and washing of cells, lung cells were stained for lineage (Ly6C, Ly6G), streptavidin, CD11b, CD64, CD11c and SiglecF.

#### Spleen

The spleens were removed from Bl6 mice and minced through a 100 μm cell strainer. After washing with staining buffer (2mM EDTA, 1% FCS in PBS) at 1,200 rpm, 4°C for 6 min, supernatant was discarded and cells were blocked with anti-CD16/32 antibodies for 10 min, before anti-F4/80-biotin antibodies were added for 25 min on ice. After washing, the cell pellet was incubated with anti-biotin beads (Miltenyi) for 15 min on ice. Cells were washed again (1,200 rpm, 4°C for 6 min) and MACS was performed with LS columns (Miltenyi). After elution and washing of cells, lung cells were stained for CD45, streptavidin, lineage (Ly6C, Ly6G, B220), CD11b, CD11c and MHCII.

#### Skin Langerhans cells

Ears were dissected and placed over 0.05% Trypsin with EDTA in PBS for 1.5-2 hours at 37°C, until the epidermis could be peeled off using forceps. The epidermis was minced and the crude tissue suspension in sorting buffer was passed through a 100 μm cell strainer and centrifuged for 6 min at 1,200 rpm and 4°C. Cell-pellets were further processed.

#### Tongue leukocytes

Tongues were extracted, minced in 500 μL of PBS with 0.2 mg of DNAse I (Roche), 2.4 mg Collagenase IV (Gibco) and 0.15 mg (60U) hyaluronidase I (Sigma) and incubated for 45 min at 37°C. 10 ml sorting buffer were added to the crude tissue suspension as it was passed through a 100 μm cell strainer and centrifuged for 6 min at 1,200 rpm and 14°C. Pellet was resuspended in 3 ml PBS, layered over 3 ml of Ficoll-Paque™ and centrifuged for 17 min at 2,000 rpm and 14°C with no acceleration nor break. The interface containing leukocytes was collected, 5 ml sorting buffer were added centrifuged for 6 min at 1,200 rpm and 4°C. Leukocyte-containing cell pellets were further processed. All tongue preparations were performed according to this protocol, if not otherwise stated.

#### Tongue Langerhans cells

To separate the epithelium from the rest of the tongue and to dislodge Langerhans cells, extracted tongues were injected with 5 U/ml of Dispase II (Sigma) in HEPES buffered saline until completely distended. They were then incubated for 15 min at 37°C and the epithelium layer was peeled off using forceps. The epithelium was minced in 500 μL of PBS with 0.2 mg of DNAse I (Roche), 4.8 mg Collagenase IV (Gibco) and 0.15 mg (60U) hyaluronidase I (Sigma) and incubated for 15 min at 37°C. 10 ml sorting buffer were added to the crude tissue suspension as it was passed through a 100μm cell strainer and centrifuged for 6 min at 1,200 rpm and 4°C. The pellet (enriched for tongue Langerhans cells) was then further processed accordingly.

### Flow cytometry and cell sorting

Cell suspensions were kept on ice. They were blocked with anti-CD16/32 (2.4G2) antibodies for 10 min and then stained for 20-25 min with antibodies against mouse CD45 (30-F11), CD45.2 (104), CD45.1 (A20), CD11b (M1/70), CD11c (N418), CD64 (X54-5/7.1), CX3CR1 (SA011F11), F4/80 (BM8), Folr2 (10/FR2), EpCam (G8.8), Gr1 (RB6-8C5), IA/IE (M5/114.15.2), B220 (Ra3-6B2), Ly6C (HK1.4), Ly6G (1A8), Lyve1 (ALY7), MerTK (2B10C42), Siglec-F (E50-2440) and TIMD4 (RMT4-54). Antibodies were purchased from BioLegend or eBioscience. The gating strategy for all macrophage populations is presented in Suppl. Fig. 3. Samples were washed in 2 ml sorting buffer and centrifuged for 6 min at 1,200 rpm and 4°C. Pellet was resuspended in sorting buffer and flow sorted using AriaI, AriaII or AriaIII (BD Biosciences, BD Diva Software) cell sorters. Flow cytometry analysis was performed on Fortessa or LSRII (BD Biosciences, BD Diva Software) and analyzed with FlowJo software v.10.7.1 (BD).

### scRNAseq

Experiment 1 (Fig. 1): Tongues were prepared as described in the material & method section. 8 adult, female Bl6 mice were used for the isolation of interstitial cells with collagenase / hyaluronidase digestion. 4 adult, female Bl6 mice were used to isolate tongue Langerhans cells with dispase digestion. Both preparations were pooled and CD45^+^ cells were analyzed with Chromium™ Single Cell 3’ Reagent Kits v3.1.

Experiment 2 (Suppl. Fig. 1a): scRNA-seq was performed on CD45^+^ tongue hematopoietic cells isolated from 10 Bl6 mice. Tongues were extracted, minced in 500 μL of PBS and digested with 0.2 mg of DNAse I (Roche) and 1mg Collagenase IV (Gibco). 10 ml sorting buffer were added and tissue suspension was passed through a 100 μm cell strainer and centrifuged for 6 min at 1,200 rpm and 14°C. Cells in the pellet were stained for anti-CD45. FACS-purified CD45^+^ cells were used for scRNA-seq. scRNA-seq was performed with the Chromium™ Single Cell 3’ Reagent Kits v2.

Experiment 3 (Fig. 4): Cell isolation of LPS-injected female Bl6 mice was performed as mentioned above. 6 mice were used for interstitial cell isolation and 4 mice for Langerhans cell extraction. Both samples were pooled and CD45^+^DAPI^−^ cells were analyzed with Chromium™ Single Cell 3’ Reagent Kits v3.1.

Experiment 4 (Fig. 5): Cell isolation of p3-p4 Bl6 mice was performed as mentioned above. 7 mice were used for interstitial cell isolation. CD45^+^DAPI^−^ cells were analyzed with Chromium™ Single Cell 3’ Reagent Kits v3.1.

Experiment 5 (Suppl. Fig. 5): Cell isolation of Irf8^−/−^ female Bl6 mice was performed as mentioned above. 5 mice were used for interstitial cell isolation. No additional Langerhans cell extraction was performed for this experiment. CD45^+^DAPI^−^ cells were analyzed with Chromium™ Single Cell 3’ Reagent Kits v3.1.

### RNA isolation and cDNA synthesis for bulk RNA-Seq

500-20,000 sorted cells were lysed with 100 μl of lysis/binding buffer (Life Technologies), snap-frozen on dry ice and stored at −80°C until further use. mRNA purification was performed with the Dynabeads™ mRNA DIRECT™ Purification Kit (Life Technologies) according to the manufacturer’s guidelines. MARS-seq barcoded RT primers were used for reverse transcription with the Affinity Script cDNA Synthesis Kit (Agilent) in a 10 μl reaction volume.

### Bulk RNA-sequencing

The MARS-seq protocol was used for bulk RNA sequencing ^49^. After reverse transcription, samples were analyzed by qPCR and samples with similar Ct values were pooled. Samples were treated with Exonuclease I (New England BioLabs (NEB)) for 30 min at 37°C and for 10 min at 80°C followed by a 1.2X AMPure XP beads (Beckman Coulter) cleanup. The second strand synthesis kit (NEB) at 16°C for 2 hours was used for cDNA synthesis followed by a 1.4X AMPure XP bead cleanup. In vitro transcription (IVT) was performed at 37°C for 13-16h with the HiScribe T7 RNA Polymerase kit (NEB). The remaining DNA was digested by Turbo DNase I (Life Technologies) treatment at 37°C for 15 min followed by a 1.2X AMPure XP bead cleanup. RNA fragmentation (Invitrogen) was performed at 70°C and the reaction was stopped after 3min with Stop buffer (Invitrogen) followed by a 2X AMPure XP bead cleanup. Ligation of the fragmented RNA to the MARS-seq adapter was performed at 22°C for 2h with T4 RNA ligase (NEB) followed by a 1.5X AMPure XP bead cleanup. A second reverse transcription reaction was performed with MARS-seq RT2 primer and the Affinity Script cDNA Synthesis Kit (Agilent Technologies) followed by 1.5X AMPure XP bead cleanup. Finally, the library was amplified using P5_Rd1 and P7_Rd2 primers and the Kapa HiFi Hotstart ready mix (Kapa Biosystems) followed by a 0.7X AMPure XP bead cleanup. Fragment size was measured using a TapeStation (Agilent Technologies) and library concentrations were measured with a Qubit fluorometer (Life Technologies). The samples were sequenced using a NextSeq 500 system (Illumina).

### Microscopy

#### Tissue preparation

Mice were deeply anesthetized with a combination of 150 mg/kg body weight pentobarbital sodium (WDT) and perfused transcardially for 3 min with ice-cold 0.9 % NaCl solution and for 5 min with 4 % paraformaldehyde (PFA) in 0.1M phosphate buffer (PB). Tongues were post-fixed overnight in 4 % PFA in 0.1M PB. They were then cryoprotected by 3 x overnight incubations in 30 % w/v sucrose in 0.1M PB, cryosectioned with a cryostat (35 μm sagittal) and mounted directly to slides for staining.

#### Immunohistochemistry

35 μm sagittal tongue sections from the tongue midline were washed for 5 min at RT in TBS (42mM Tris HCl, 8mM Tris Base, 154mM NaCl; pH 7.4). Sections were blocked in 20 % normal donkey serum (NDS) in TBS-T for 1 hour at RT and incubated overnight at 4°C in primary antibody diluted in TBS-T + 2 % NDS/NGS. Primary antibodies against CD31 (Millipore MAB13982; 1:200), CD68 (Bioloegend 137002; 1:100), GFP (AbCam; ab13970; 1:200), Lyve-1 (ReliaTech 103-PA50AG; 1:500), Podoplanin (Biolegend 156202; 1:50), Tuj1 (AbCam ab18207; 1:200) were used. Sections were then washed 3 x 30 min in TBST and incubated overnight at 4°C with appropriate secondary antibodies raised in goat and conjugated to Alexa Fluor dyes (Invitrogen) were diluted 1:500 or 1:1000 in TBS-T + 2 % NDS/NGS. Sections were washed 3 x 30 min in TBST and nuclei were stained with DAPI at a 0.5 mg/ml in TBS. After washing, sections were mounted with FluorSave reagent (Calbiochem).

#### Imaging and quantification

Imaging was performed using a Leica SP5 TCE. Four regions (R1-R4; Fig. 3e) from three consecutive sagittal 35 μm slices at the tongue midline were imaged at 775.76 x 734.76 μm dimensions and over a depth of 30 μm with 1.5 μm z-stack intervals. All image processing was done with Fiji ^50^ and quantifications were performed with Imaris Microscopy Image Analysis Software (Bitplane). Regions of interest (ROI: muscle or lamina propria) were manually drawn onto each z-stack image and the software’s built-in surface module extrapolated the volume of the respective ROI. Then, the built-in surface detection algorithm was used to identify cells (LYVE1^+^ or *Cx3cr1*-GFP^+^) in each ROI, so that we could calculate number of cells and mean cell volumes per ROI volume.

### Lipopolysaccharide induced systemic inflammation

Mice were injected intraperitoneally with 1 mg/kg LPS (E. coli 0111:B4) in 200 μl PBS 6 hours prior to sacrifice.

### Cell proliferation assay (EdU)

The EdU Click 488 Kit from BaseClick were used. In brief, 0.5 mg/g of EDU in PBS was injected intraperitoneally to adult mice or subcutaneously to pups 15 hours before to sacrifice. Single cell suspensions were obtained from tongues (see relevant section) and cells were processed according to the manufacturer’s instructions.

### Sequencing Data Analysis

#### Bulk sequencing data analysis

Using the fastq files, the reads were deduplicated based on their UMIs. Subsequently, STAR (version 2.5.3a) was applied for the alignment of reads to the mouse genome (mm9). Quantification of reads by htseq-count (version 1.0) yielded the input expression matrix for DESeq2 (version 1.26.0), used to identify differentially expressed genes for each group. More specifically, each group was compared against all other groups to obtain DE marker genes.

### Single cell sequencing data analysis

#### Preprocessing and integration

The data was sequenced with 10x Genomics (version v3.1) For alignment to mm10 and quantification Cell Ranger Single Cell Software was used. For additional preprocessing and downstream analysis Seurat (v4) was applied. First, based on the UMI counts, cells were filtered based on the number of detected genes and the proportion of mitochondrial gene counts. All cells with more than 10% mitochondrial gene count were removed. For the number of detected genes, a sample specific filtering value was applied, for experiment 2 (10x Chromium v2) the accepted range was between 300 and 2000, for the remaining samples a minimum of 500 was required while the maximum cutoff ranged from 5000 (experiment 1) over 5500 (experiment 3) to 6000 (experiment 4). The data was normalized and the 4000 most valiable genes were selected. Subsequently, the data was analyzed together by applying Seurats integration function (FindIntegrationAnchors) and making use of the pre-computed anchors (FindIntegrationAnchors), i.e. genes used to map the cells from different samples. Analogously, experiment 5 (*Irf8*-deficient cells) and experiment 1 were integrated. For the IRF8 sample, the range of detected genes for included cells was set to 500-5500 and cells with more than 10% mitochondrial gene count were removed.

#### Dimension Reduction, unsupervised clustering and marker detection

A principal component analysis provided the basis for the computation of a UMAP and an unsupervised clustering of the cells. The resolution parameter for FindClusters was set to 0.42 (0.25 for experiment 5). Conserved cluster markers for all samples were computed using FindConservedMarkers with the detection method set to ’MAST’. Genes differentially expressed between clusters of different samples were detected analogously using FindClusters.

#### Cell cluster annotation, trajectory inference and signature enrichment

To assign cell types to the clusters identified, we applied SingleR (version 1.6.1; ^14^), using the ImmGen database as annotation resource. Slingshot (version 2.0.0; ^28^) was applied for trajectory inference. To this end, we provided slingshot with the newborn-specific cluster as starting point. To shed more light on the macrophagic versus dendritic nature of the cells in the central clusters, we computed the enrichment of macrophage-specific as well as dendritic gene sets by use of GSVA (version 1.40.1; ^15^).

## Data and code availability

Data that were generated within this study have been deposited in Gene Expression Omnibus (GEO) with the accession code GSEXXXX.

## Acknowledgement

We would like to thank Jermaine Voß for excellent technical support, as well as the MDC animal facility, especially Juliette Bergemann, the MDC FACS core unit and Dr. Hans-Peter Rahn in particular, and the MDC genomic core facility, especially Caroline Braeuning. We thank Dr. Anca Margineanu, Dr. Sandra Cristina Carneiro Raimundo and the Advanced Light Microscopy Technology Platform of the MDC for the general and technical support. We are also grateful to Dr. Christoph Klose for advice on lymphocyte subset identification. A.M is a Heisenberg fellow supported by the DFG (MI1328).

## Author contribution

E.M.L., L-K.W. and D.D. performed experiments and analysis. K.Z. performed bioinformatic analysis. C.F. and T.C. helped with scRNA-seq experiments. A-H.H, S.Y., S.J., W.S. and S.D. provided critical mouse strains, reagents and intellectual input. A.L. provided resources, editing and lab space. A.M. financed, designed and supervised the study. A.M. and E.M.L wrote the manuscript.

---

**Supplementary Figure 1:**
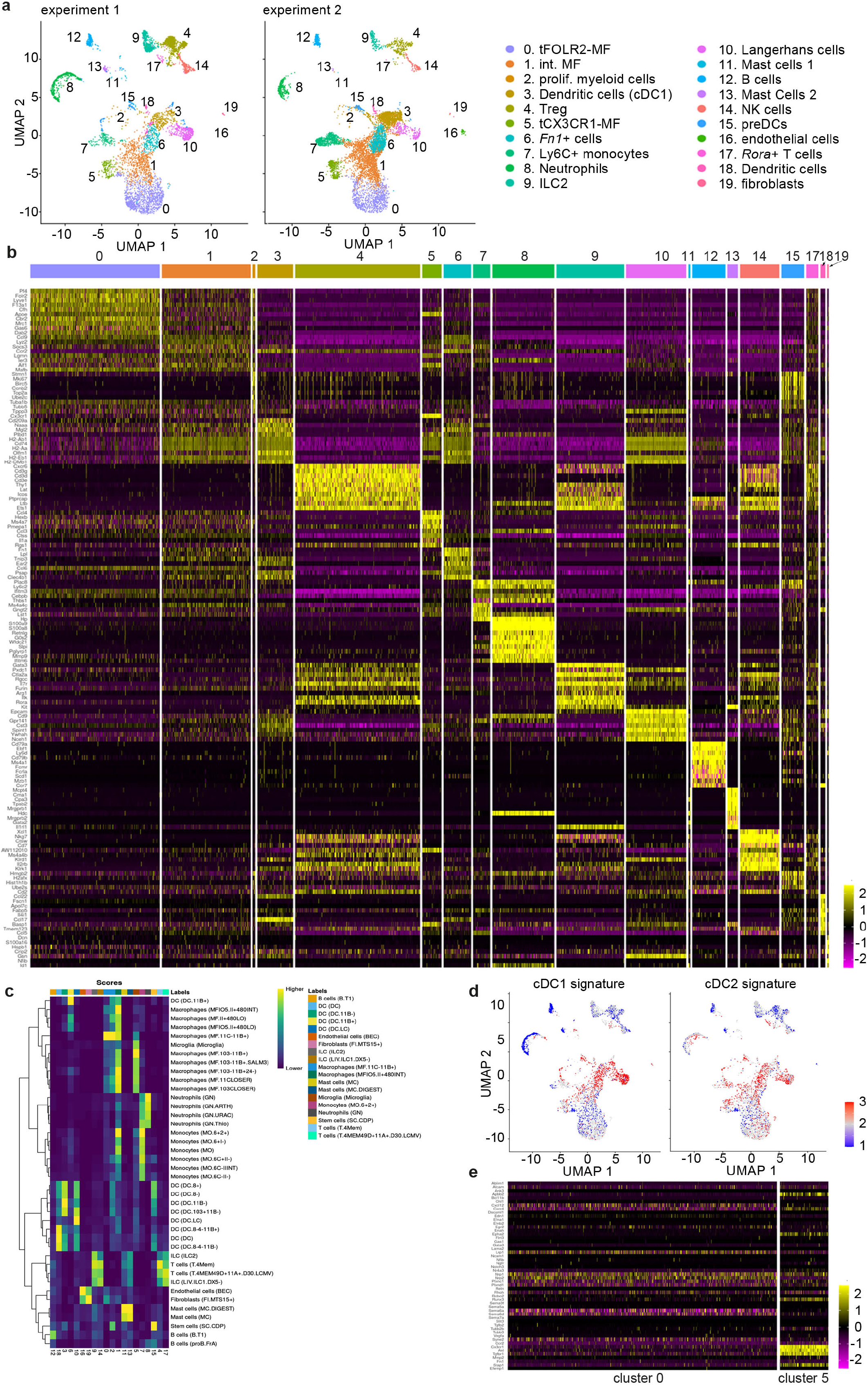
Characterization of adult tongue leukocytes (**a**) Biological and technical replication (experiment 2, right graph) of the scRNA-seq experiment depicted in Fig. 1 (experiment 1, left graph). For experiment 2, CD45^+^ tongue leukocytes were isolated from female mouse tongues and 8151 cells were sequenced using the Chromium Single Cell 3’ Reagent Kits v2. Experiment 2 data was integrated with the existing adult data of experiment 1 to allow population comparison. (**b**) Heatmap representing the top 10 marker genes for all detected clusters. Data is derived from experiment 1 shown in Fig. 1. (**c**) SingleR results for the annotation of cell clusters. Note that cluster 6 cells showed similarities to CD11b^+^ DCs from Immgen (Immgen.org). (**d**) GSVA analysis of the cDC1 and cDC2 signature genes as defined in ^17^. Red indicates cells with higher enrichment scores for the signature genes compared to blue. (**e**) Heatmap of genes involved in axon guidance. Shown are the normalized read counts in tFOLR2-MF (cluster 0) and tCX3CR1-MF (cluster 5).

**Supplementary Figure 2:**
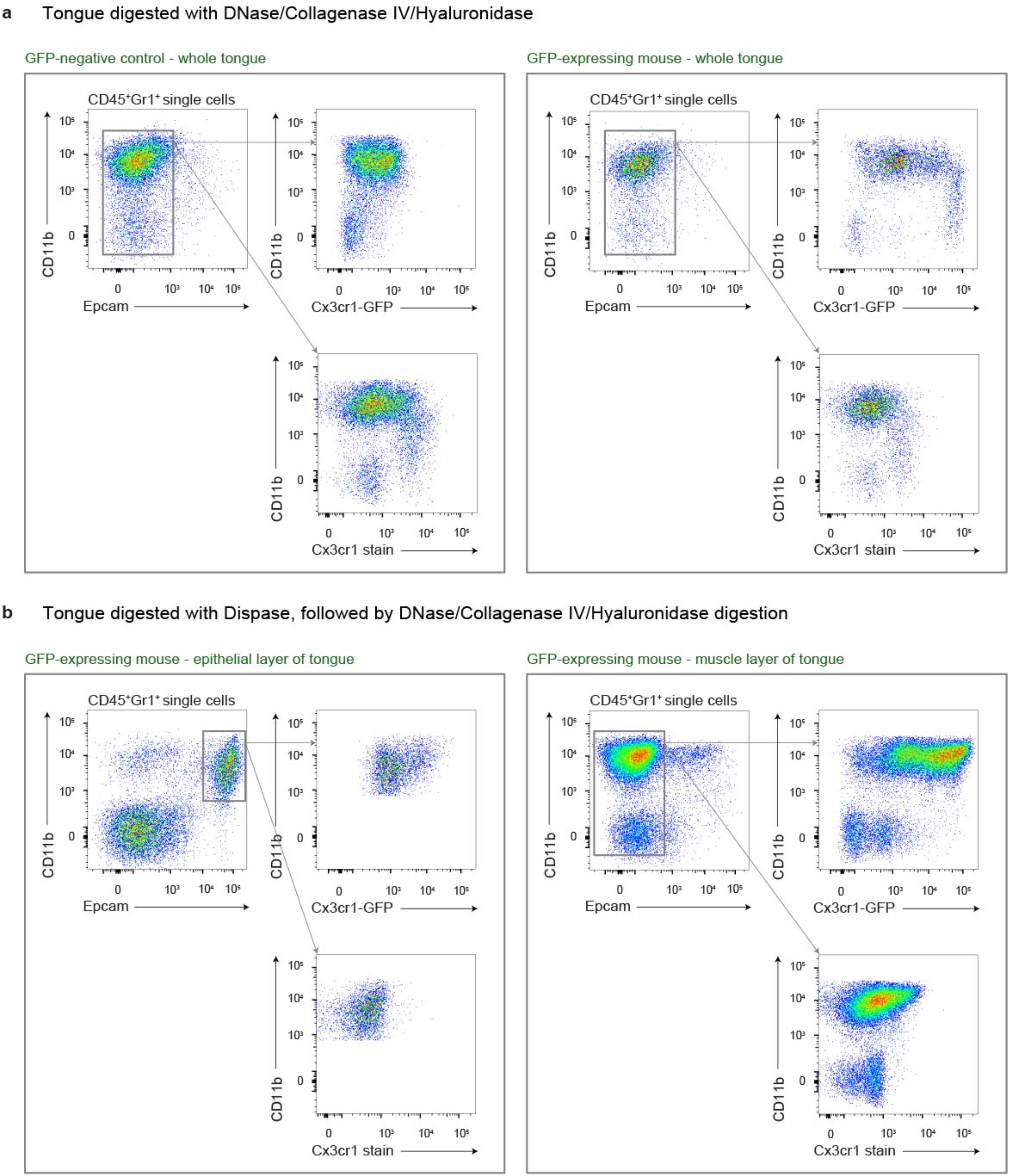
Different isolation methods for the identification of tongue leukocytes (**a**) Digestion of tongues from PBS perfused mice with collagenase IV, hyaluronidase and DNase. Shown are examples for Bl6 mice (left) and *Cx3cr1*^Gfp/+^ mice (right). Note the almost complete absence of CD11b^+^ Epcam^+^ Langerhans cells. (**b**) After *Cx3cr1*^Gfp/+^ tongues were first digested with dispase, the epithelial layer was peeled off and subsequently digested with collagenase IV, hyaluronidase and DNase. With this preparation protocol, CD11b^+^ Epcam^+^ Langerhans cells could be detected in the epithelial layer (left), but not in the muscle layer (right).

**Supplementary Figure 3:**
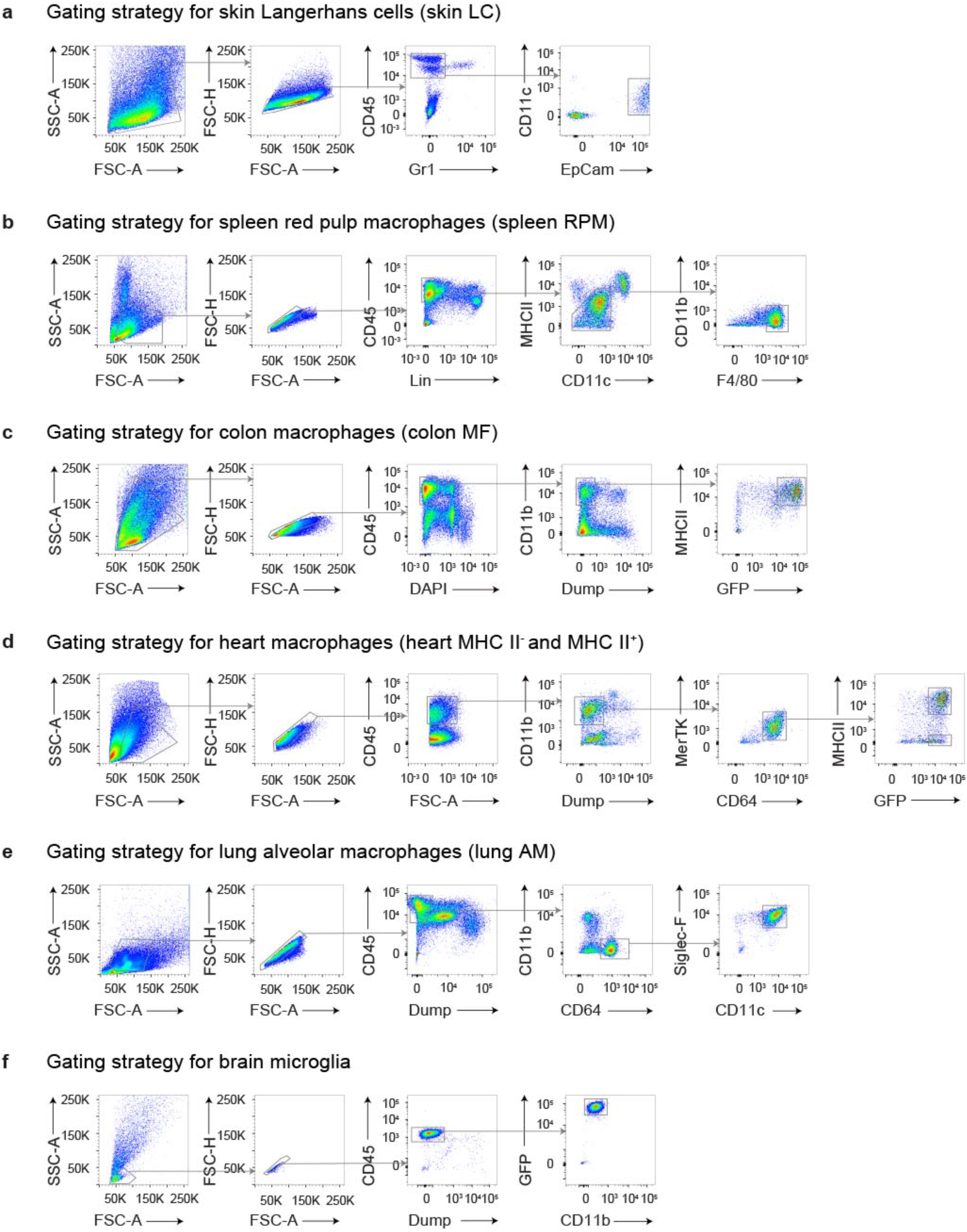
Gating strategy for the isolation of tissue resident macrophages Shown are the gating strategies that allow the identification of (**a**) Langerhans cells in the skin, (**b**) splenic macrophages, (**c**) *Cx3cr1*^+^ colonic macrophages, (**d**) MHCII^+^ and MHCII^−^ *Cx3cr1*^+^ heart macrophages, (**e**) alveolar macrophages and (**f**) microglia from the brain.

**Supplementary Figure 4:**
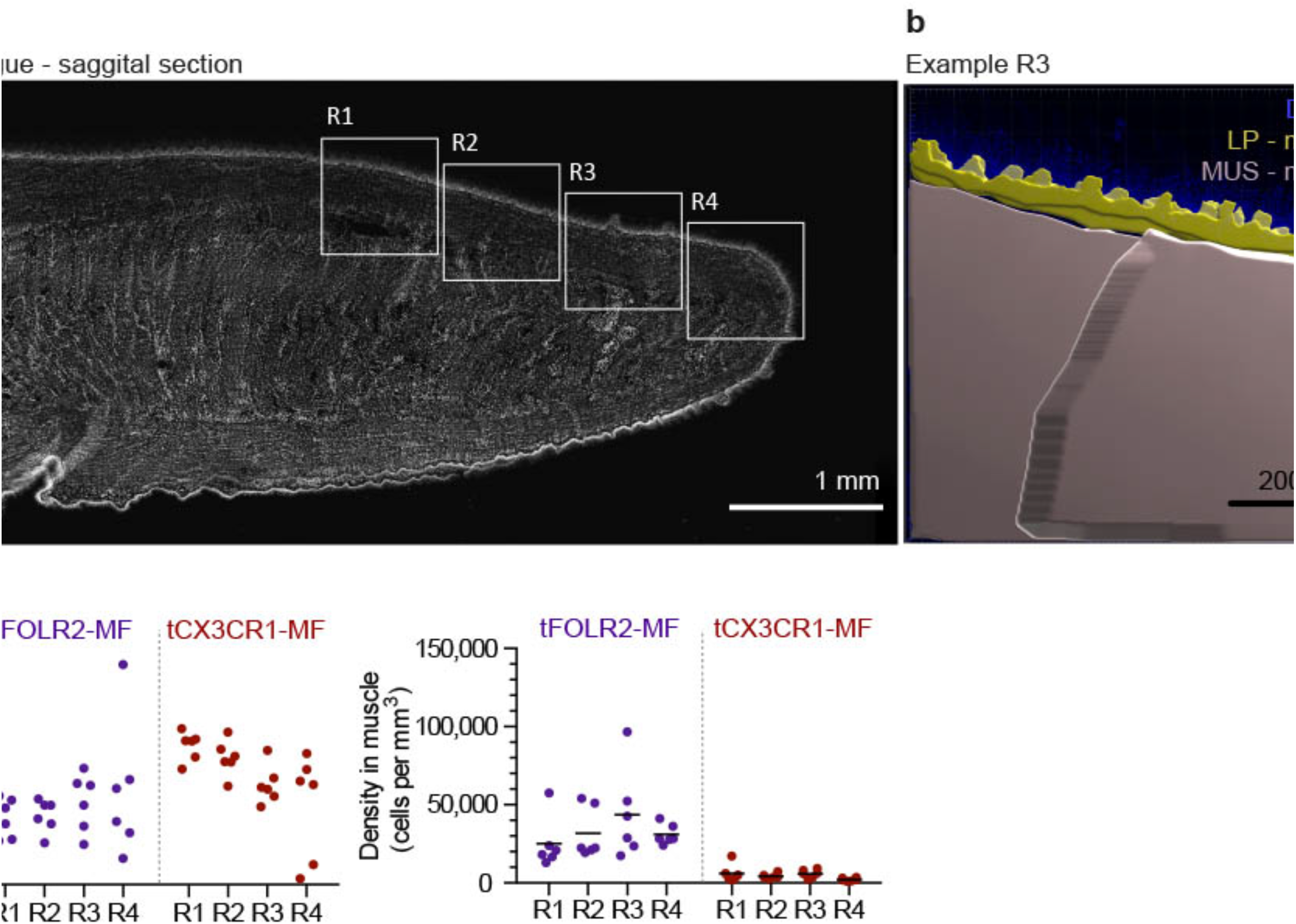
Quantification of tongue macrophages (**a**) Panoramic image of an adult tongue counterstained with DAPI (white). The insets represent different areas of the tongue that were quantified. (**b**) Imaris-based mask for the identification of epithelium, lamina propria and muscle. Analysis of cell density was performed within these areas. (**c**) Macrophage subset cell counts in the respective regions of the tongue as represented in (a). Shown is the density of cells / mm^3^ in the lamina propria (left) and in the muscle (right). Each dot represents one independent animal.

**Supplementary Figure 5:**
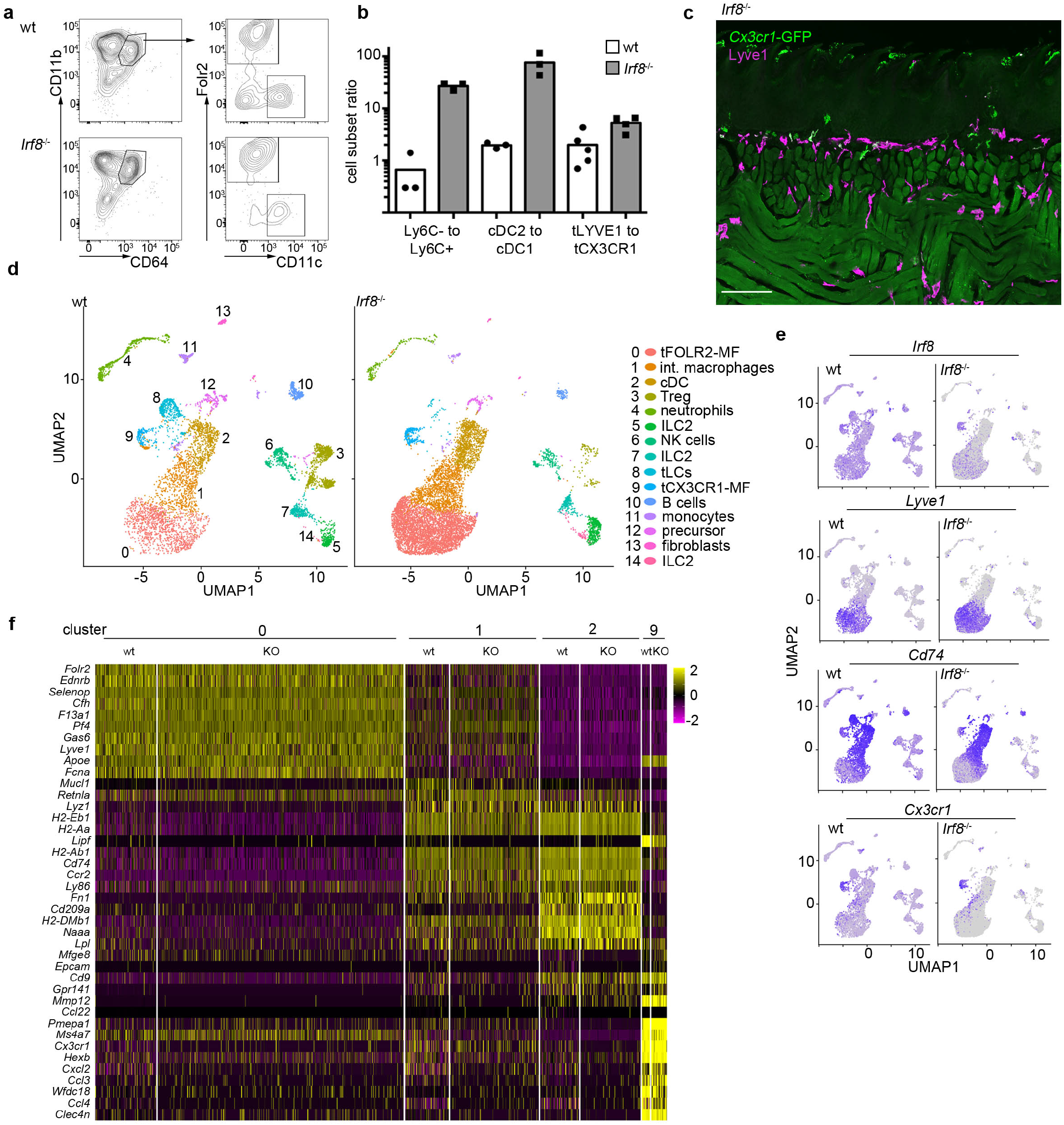
*Irf8*-deficiency does not affect the tongue leukocyte composition (**a**) Flow cytometry analysis of *Irf8*-deficient tongue leukocytes and their respective controls. The cells were pre-gated as CD11b^+^ CD64^+^. (**b**) Ratio analysis of Ly6C^−^ monocytes / Ly6C^+^ monocytes, splenic cDC2 / cDC1 cells and tFOLR2-MF / tCX3CR1-MF isolated from WT (n = 3-5) and *Irf8*-deficient mice (n = 3-4). Each dot represents one animal. The experiment was repeated twice. (**c**) Histological analysis of an adult *Cx3cr1*-GFP Irf8^−/−^ tongue reveals the presence of GFP^+^ tCX3CR1-MF in the lamina propria. Sections were stained with anti-GFP (green) and anti-LYVE1 (violet) antibodies. (**d**) 9047 CD45^+^ cells were profiled from adult *Irf8*-deficient mice (pool of n=5 mice) by scRNA-seq. Shown are UMAP dimension reductions of WT (left) and Irf8-deficient CD45^+^ tongue cells (right). (**e**) Gene expression examples in wt and *Irf8*-deficient cells. Note that *Irf8* transcripts can be detected on mRNA level in *Irf8*-deficient mice since only exon 2 of the *Irf8* gene is deleted in the Jackson strain 018298. (**f**) Heatmap of marker gene expression for clusters 0, 1, 2 and 9 in wildtype and Irf8-deficient cells.

Supplementary data 1: Marker genes for the 19 identified clusters in the scRNA-seq experiments 1-4 together with the respective test statistics. Each cluster can be found in separated tabs. As test method we used MAST, the log2fc threshold was set to 0.25 and a Bonferroni mutiple testing correction was applied. We only considered genes, that were expressed in at least 20% of the cells in at least one of the groups. Note that cluster 11 is not represented due to low cell numbers and cluster 16 could only be detected in experiment 3 (LPS). These data sets belong to Fig. 1, 4 and 6.

Supplementary data 2: The file contains the average expression value for each cluster based on the normalized counts of the cells in the respective sample. Additionally, a column for each cluster indicates, whether a gene is a conserved marker for this cluster (see Suppl. Data 1). This data set belongs to Fig. 1.

Supplementary data 3: Core gene signatures of cDC (tab 1), macrophages (tab 2), cDC1 (tab 3) and cDC2 (tab 4). The gene lists were derived from ^16,17^. This data set belongs to Fig. 1.

Suppl. data 4: Represented are the full GO lists for the myeloid clusters 0, 1, 3, 5, 6, 7 and 10. This data set belongs to Fig. 1.

Suppl. data 5: This table includes the full bulk RNA-seq read counts for all isolated tissue resident macrophage populations that are shown in Fig. 2. Each subset is represented in a separate tab.

Suppl. data 6: Listed are the upregulated genes that could be extracted from the bulk RNA-seq data. These genes were used for GO annotations represented in Fig. 2.

Suppl. data 7: Listed are all significant differential expressed genes between untreated and LPS-treated tFOLR2-MF (tab1) and tCX3CR1-MF (tab2).

Suppl. data 8: This html file shows the 3D representation of the postnatal day 3 UMAP represented in Fig. 6

